# A Quantitative Modelling Approach for DNA Repair on a Population Scale

**DOI:** 10.1101/2022.03.29.486283

**Authors:** Leo Zeitler, Cyril Denby Wilkes, Arach Goldar, Julie Soutourina

## Abstract

Despite intensive research of DNA repair after UV in eukaryotes, a framework to quantitatively describe the dynamics *in vivo* is still lacking. We developed a new data-driven approach to analyse CPD repair kinetics over time in *Saccharomyces cerevisiae*. In contrast to other studies that consider sequencing signals as an average behaviour, we introduce a hidden axis representing independent cells where loci can transition from *damaged* to *repaired*. This permits the application of the Kolmogorov-Johnson–Mehl–Avrami model to find a region-specific and continuous representation of the entire temporal process. We correlated the parameters via a *k*-nearest neighbour approach to a variety of genomic features, including transcription rate and nucleosome density. The clearest link was found for the gene size, which has been unreported for budding yeast to our knowledge. The framework hence allows a comprehensive analysis of nuclear processes on a population scale.

**Author Summary:** As DNA encodes our very identity, it has been subject to a plethora of studies over the last century. The advent of new technologies that permit rapid sequencing of large DNA and RNA samples opened doors to before unknown mechanisms and interactions on a genomic scale. This led to an in-depth analysis of several nuclear processes, including transcription of genes and lesion repair. However, the applied protocols do mostly not allow a high temporal resolution. Quite the contrary, the experiments yield often only some few data signals over several hours. Missing dynamics between time points are chiefly ignored, implicitly assuming that they straightforwardly transition from one to another. Here, we show that such an understanding can be flawed. We use the repair process of UV-induced DNA damage as an example to present a quantitative analysis framework that permits the representation of the entire temporal process with only three parameters. We subsequently describe how they can be linked to other heterogeneous data sets. Consequently, we evaluate a correlation to the whole kinetic process rather than to a single time point. Although the approach is exemplified using DNA repair, it can be readily applied to any other mechanism and sequencing data that represents a state transition between two states, such as *damaged* and *repaired*.

## Introduction

As DNA represents the hereditary unit of life, maintaining its integrity is vital for every organism’s survival. A large variety of different genotoxic factors have the potential to damage the molecular structure of DNA. Among others, it has been shown that UV light induces Cyclobutane Pyrimidine Dimers (CPDs). Nucleotide Excision Repair (NER) is an evolutionary conserved repair mechanism in *Saccharomyces cerevisiae* that can remove a broad range of damage, including CPDs ([25]). NER is conventionally divided into two subpathways. The first recognition mechanism is named Global-Genome Repair (GGR). DNA damage is recognised directly by the Rad4-Rad24-Rad33 complex. There is evidence that protein loading is promoted through interactions with chromatin remodellers that change the nucleosome density or distribution ([15, 36]). The second pathway is restricted to actively transcribed regions; hence the name Transcription-Coupled Repair (TCR). Expressed genes exhibit quicker repair than silent downstream regions ([6]). This promoted the assumption that TCR is more efficient than GGR, although constrained to the transcribed strand (TS) ([30, 39]). TCR is initiated by lesion-blocked RNA polymerase II (Pol II) which cannot continue elongation ([9]). Thus, a potential link of TCR to transcription rate has been indicated by several studies ([26, 28]). After recognition, TCR and GGR use the same incision and nucleotide replacement mechanism. DNA is incised to either side of the lesion leaving an approximately 30-nucleotide gap, which is subsequently replaced and ligated (for a comprehensive description and analysis, see the review by [7]).

Our understanding of such processes in living cells has been largely enhanced by Next Generation Sequencing (NGS). It allows the identification of enriched loci of a selected property on a genome-wide scale. Among others, it has been applied to investigate the *in vivo* CPD repair mechanisms through analysing temporal changes of the damage distribution. [28] obtained high-resolution CPD-seq data that are often used as a benchmark reference (see for example [26]). Their analysis indicates that single nucleosomes and DNA-bound transcription factors have an impact on the CPD formation. Moreover, they point out that repair is seemingly influenced by the CPD position with respect to the nucleosomal dyad as well as the transcription rate of genes. Another major contribution has been done by [26]. Their protocol for excision repair sequencing (XR-seq) revealed strong TCR at early time points which is followed by repair in non-transcribed regions. Furthermore, [46] and [41] utilised CPD data to compute repair rates in different areas, which indicated that the process is highly organised into genomic regions. By using GGR-deficient strains, they show that repair is changing globally when the subpathway is repressed. This is compared to the distribution of repair proteins and histone modifications.

Although all of these studies analyse the sequencing data in detail, it is important to highlight that they consider the CPD signal as being representative for an average cell. Such a notion—and hence the possibility to make conclusions about repair in single cells—presumes that they do not influence each other’s repair processes in a culture. Otherwise, any conclusion would be only population-specific. In order to explicitly take into account these independent kinetics, we assume that the CPD-seq signal corresponds to a collapsed data grid with two dimensions, i.e. the position along the genome and the cells in a population (see top row in Fig 1). Every position in the grid can be in either of two states, *damaged* or *repaired*. During ongoing repair, it is presumed to observe an irreversible transition from a *damaged* to a *repaired* state. The signal expansion causes the DNA lesions to be scattered over the entire population. We assume that this results in the actual distribution of lesions within a single cell (e.g. a row in the grid) to be scarce. Thus, considering the expected spatial distance between two lesions, we hypothesise non-influencing repair. It follows that rows and columns in the data grid are independent of each other. Repaired regions can be consolidated to patterns, which subsequently grow over time (see bottom row of Fig 1). This strongly resembles of the crystallisation and growth kinetics as described in detail by [24], [20] and [2–4] (KJMA model). The one-dimensional version of KJMA theory has been successfully applied in a biological context to analyse the dynamics of DNA replication in eukaryotes ([22]). Here, we use a two-dimensional version to assess the DNA repair dynamics on a population scale.

**Figure 1.**
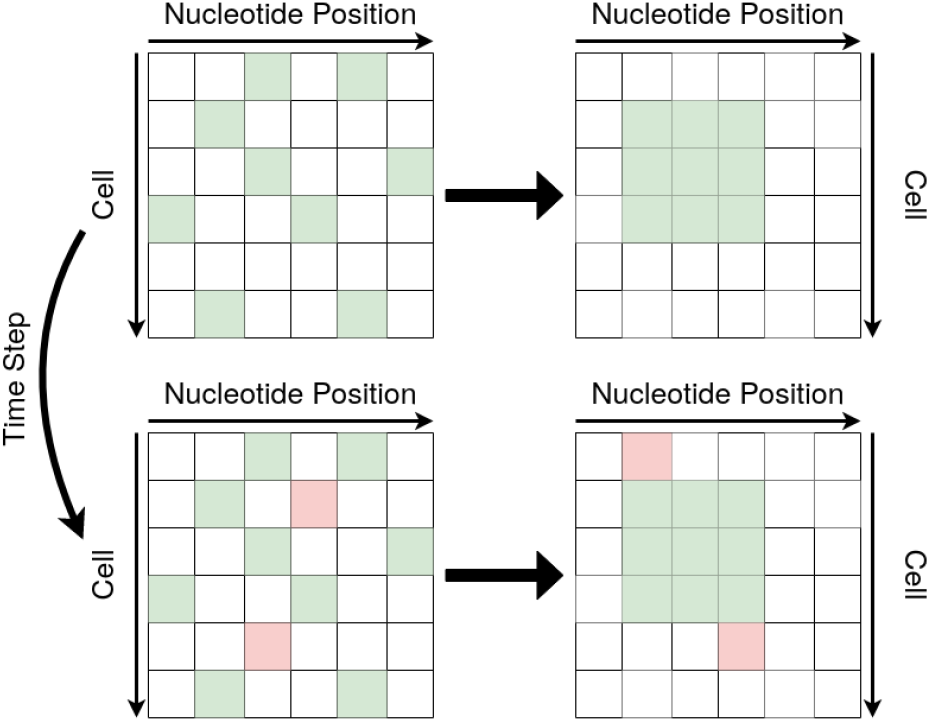
Re-arranging repaired positions to create patterns. Repaired positions per cell and per locus distributed on a grid (green). When assuming repair is independent within and between cells, positions can be reordered to create patterns. This is done for every genomic region, and it is thus context-specific (e.g. transcript vs. intergenic region). Newly repaired lesions (red) at later time steps are added to existing patterns.

In this study, we present a computational framework for analysing DNA repair kinetics. To our knowledge, it is the first time that independent cells are explicitly taken into account when assessing sequencing data over time. Most importantly, the theory allows a state description without presuming a particular molecular mechanism. In the following, we firstly present how the KJMA model can be adapted and applied to CPD repair. We introduce an abstract repair space that allows us to link the physical model with the biological process. As a consequence, we can make predictions about the expected temporal evolution of sequencing signals. We determine descriptive parameters for a large number of DNA regions. The theoretical contribution is complemented by successfully linking the derived parameters—which now represent the temporal process rather than a static distribution—with other genomic properties. This includes heterogeneous data from various studies, inter alia nucleosome density, transcription rate and transcriptional unit length. The significance of all correlations are computationally verified using a non-parametric *k*-nearest neighbour (*k*NN) approach. As supported by other findings, our results show an interrelationship of TCR with transcription rate ([26, 28]). Furthermore, we could find a link to nucleosome density, especially in intergenic regions. A correlation with nucleosomal organisation has also been proposed by [46, 47] and [41]. The clearest association, however, was found between repair parameters and gene size. To our knowledge, this is unreported for budding yeast. The presented quantitative framework is hence able to recover known influencing factors as well as predict new interrelationships. This demonstrates the generality and validity of our method, which can be readily extended for assessing the signal evolution in various chosen contexts. Furthermore, it is computationally lightweight due to its simplicity. The source code is available on GitHub: https://github.com/leoTiez/jmak.

## Results

### Motivating the Application of the KJMA Model

The exact kinetics that drive CPD repair *in vivo* remain largely unknown. Fortunately, available sequencing data permits the analysis of nuclear factors that are involved in the lesion removal process. Any further downstream analysis needs to account generally for two major issues. Firstly, we assume a considerable amount of noise due to the cellular process, but also because of the data acquisition itself. Secondly, probing time points of sequencing data are commonly coarsely distributed, with often only three to four data values over several hours ([26, 28, 41, 46]). It is hence necessary to find a reasonable functional representation of DNA repair kinetics to study the temporal process. We assume the following:

1. DNA repair is independent between cells. Cell *A* does not influence repair in cell *B*.
2. DNA repair is independent within the same cell. Locus *x* does not influence repair of locus *y*.

In the following, we converted the CPD data to a repair distribution unless otherwise stated (see Eq 4). Considering the aforementioned two-dimensional grid, the two assumptions above permit us to re-order repair positions to patterns. Despite assuming independence of repair kinetics, we assume that they are heterogeneous along the genome. Multiple studies showed that the TS of some genes is more efficiently repaired than the non-transcribed strand (NTS) ([26, 28, 41]). Even within transcribed regions, the beginning and centre of a gene exhibits seemingly a stronger CPD decrease than the end ([28]). Therefore, we restrict the re-grouping to stay within areas of interest which are assumed to behave homogeneously (Fig 1, top). In other words, we aim to find a functional description for each considered genomic region. Instead of describing the total (or relative) changes of the one-dimensional CPD-seq graph, this interpretation allows the analysis of a pattern expansion (exemplified in Fig 1, bottom). The KJMA model provides a well established framework for this analysis.

### Modelling Repair Kinetics as a Phase Transition

The KJMA model describes the fraction of transformed particles in a phase shift of solids at constant temperature (see Methods and Materials for more details, in particular Eq 11). We adapted it to explain the evolution of CPD repair. The fraction of transformed particles *f* (*t*) is re-defined, by analogy, to be the fraction of repaired lesions. Due to the independence assumptions, the expansion of re-ordered patterns can be explained by an expansion speed *G*. In the following, we assume *G* to be constant in all directions. *f* (*t*) changes as a function of time with respect to *G* as well as the amount of remaining CPDs. The original KJMA model assumes that all particles can be transformed from one phase to the other. To incorporate a cell-to-cell variability to repair UV-induced DNA damage, we introduce the parameter *θ*. It represents the maximum observed fraction of CPDs that are repaired in a population. This is expressed by the following equation:

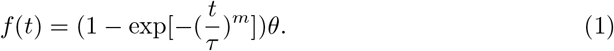

*m* is the Avrami exponent and characterises the geometry of the area covered by repaired positions after their aggregation. For example, if *m* − 1 = 2, the area corresponds to regular disks in a two-dimensional space. Irregular forms can be expressed with non-integer values ([5]). In the following, we also refer to it as a *shape* parameter. However, a direct comparison of a physical shape with a virtual pattern might be difficult to imagine. We therefore advocate another interpretation. *m* can be understood to express time dependence of the repair process (compare with Fig 2). A similar notion has been also proposed in the physical context ([8]). Furthermore, *τ* is the characteristic time, whereas 1*/τ* is the turnover or conversion rate from a *damaged* to a *repaired* state. An example of Eq 1 for different *τ* and *m* is given in Fig 2.

**Figure 2.**
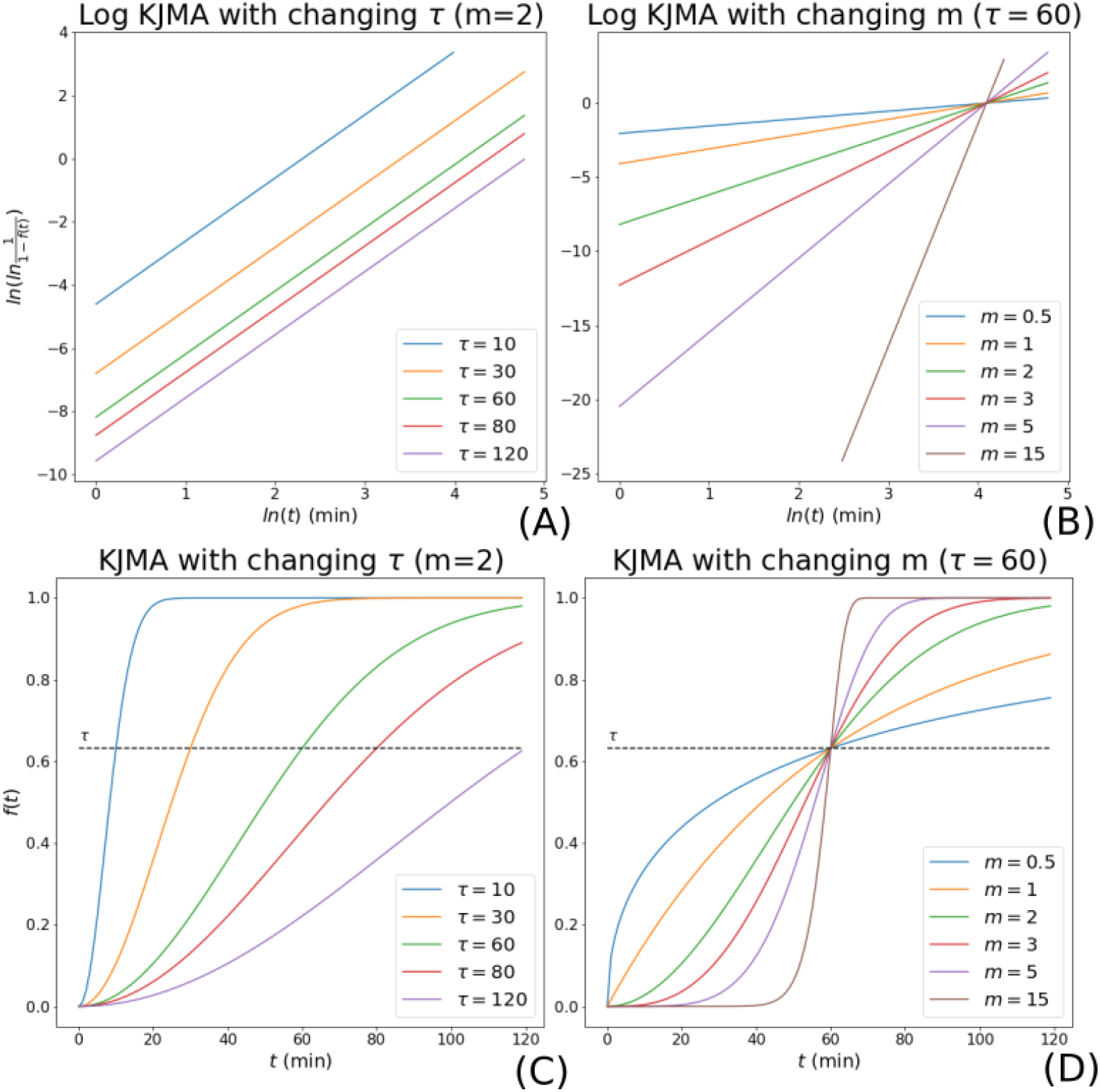
Examples of the KJMA model. (A) The KJMA model changes with different characteristic times *τ*. The larger *τ*, the more time is necessary for the process to reach 1 − 1*/e* (black dashed line). (B) Influence of shape parameter *m* on the repair dynamics. The KJMA model can be approximated by a step function with large *m*. (C, D) The repair fraction evolution transformed with respect to Eq 2.

Eq 1 explains CPD repair as an S-shaped transformation over time (see example in Fig 2). It should be noted that the process has a defined starting point at *t* = 0.

By applying the natural logarithm on both sides twice, we obtain

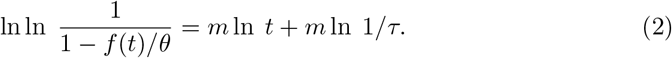

Note that the expression is continuous over *ln t*. Given the data points for repair and by assuming a value for *θ*, we can find *m* and 1*/τ* by solving the linear regression problem defined in Eq 2 (compare with bottom plots in Fig 2). *θ* can be determined through a systematic parameter search. We chose the *θ*-value that can best describe the repair data together with the corresponding parameters *m* and 1*/τ*. This is determined by maximising the adjusted *R*^2^. It represents the variance in the data that can be explained through the model and can be interpreted as goodness of fit.

This understanding of the process leads to two important conclusions. Firstly, the temporal changes in our observable—i.e. the CPD-seq data—can be non-linearly dependent on time. This becomes clear when taking the derivative and re-arranging Eq 1. We obtain

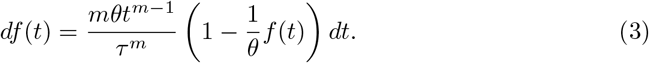

When *m* − 1 ≠ 1, *df* (*t*) changes non-linearly with time. This plays a crucial role for analysing the repair kinetics, e.g. when making conclusions about the expected repair time for a gene. Whilst it is no new finding by itself that repair is changing non-linearly in time, it is usually neglected in the actual data analysis. As a consequence, it was only permitted to compare directly the available time points without interpolating between them. Secondly, the value of *m* provides some valuable information about the repair process. As we initially postulated *G* to be constant over time in all directions on a two-dimensional grid, we would expect *m* − 1 = 2. However, Eq 2 does not impose such a constraint, and any *m* ≥ 0 could be theoretically derived based on the data. An *m*-value that sufficiently deviates from *m* − 1 = 2 indicates non-constant repair. As a result, we implicitly allow *G* to be changing over time in a two-dimensional space.

### Explaining CPD Repair Kinetics

CPD data was taken from [28] and transformed to represent relative repair with respect to the initial amount of induced CPDs (see Eq 4). Next to the traditional differentiation between gene (TS and NTS) and intergenic areas, we also introduced the notion of TCR regions. They are defined as transcripts that exhibit strong repair within the first 20 minutes, i.e. ≥20% of CPDs are removed (S1 Appendix, S1 Fig). All other genomic segments are non-TCR areas. We hence distinguish between two setups: the *gene* configuration, which consists of the genic TS and NTS as well as plus and minus strand of intergenic regions; and the *TCR* setup, which is composed of the TS and NTS TCR regions as well as non-TCR areas (see Fig 3). For a more detailed analysis of TCR regions, we further partitioned the TS and NTS into beginning, centre, and end.

**Figure 3.**
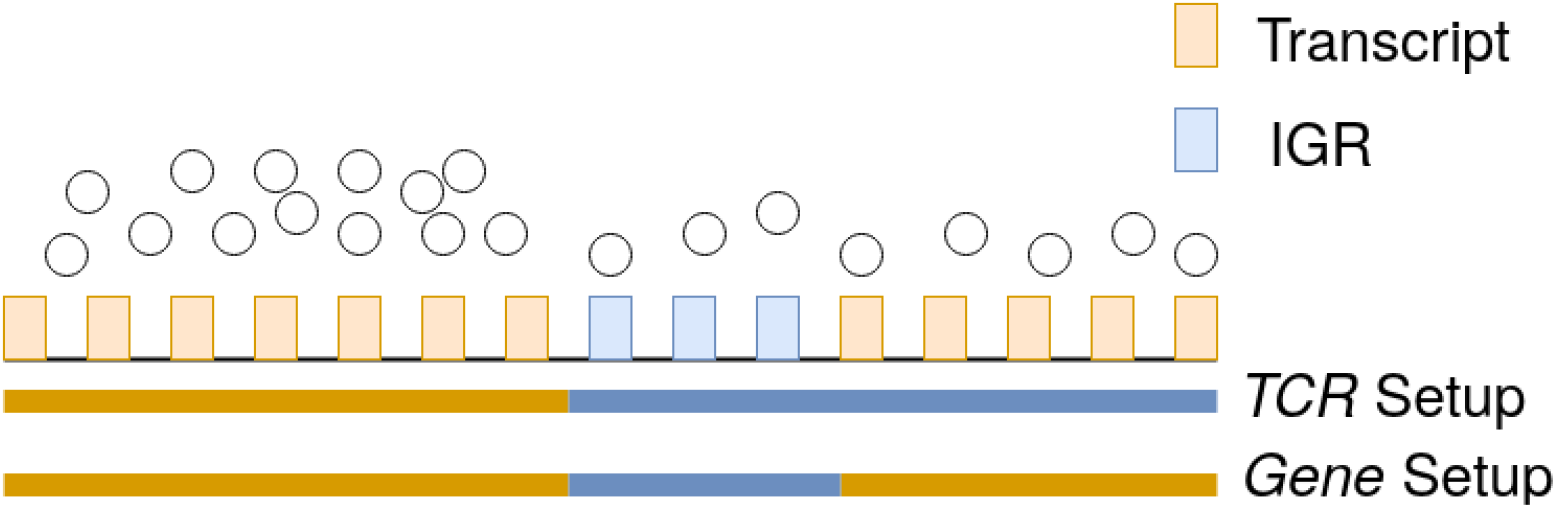
Scheme of the segmentation setup. The circles represent the number of cells with ongoing repair in the region. The positions that are marked in orange give the transcribed areas; those in blue are non-transcribed. In the *TCR* setup, only the first gene is considered as TCR area that shows more efficient repair than intergenic regions within the first 20 minutes after UV irradiation. All other parts are labelled as non-TCR region. Therefore, it spans from the end of the first gene to the end of the second. The *gene* configuration partitions the genome into the traditional notion of transcribed and intergenic regions. The gene positions were determined by [32].

Eq 1 was used to fit the parameters to the data. An example is given in S2 Fig. It is important to compare the model predictions with independently probed data to assess its validity. XR sequencing provides a snapshot of currently ongoing repair in the cell culture. We therefore suppose that it correlates with the derivative of the repair evolution (see Eq 3). Data were taken from [26]. XR-seq signals at 5, 20 and 60 minutes after irradiation were averaged with respect to the *gene* or *TCR* configuration. Due to the fact that Eq 1 and 3 are temporally continuous when *t* ≥ 0, it is straightforward to make predictions for the required time points. It hence increases comparability between different experiments. This is, however, not true for the available relative repair data, as it is dependent on the CPD distributions measured at 20, 60 and 120 minutes. We calculate the *relative repair rate* data by taking the difference to the preceding time point. This was subsequently re-scaled to represent the same time step. The model prediction and the *relative repair rate* data were plotted with respect to the XR-seq values. We opted for applying the distance correlation (DC) as a trend measurement, which captures also non-linear trends. Strikingly, the estimated *relative repair rates* correlate even better (DC=0.441) than the actual data in the *TCR* setup (DC=0.209) (compare Fig 4 (A) with 4 (B)). Moreover, the DC is fairly strong for the model predictions. The trajectories are exemplified in S3 Fig. We hence surmise that Eq 1 reasonably represents lesion removal dynamics in yeast. It should be mentioned that the DC is much lower for the *gene* configuration (DC=0.241) (S4 Fig (A)). Surprisingly, this is also true when comparing XR-seq with the *relative repair rate* data (DC=0.231) (S4 Fig (B)). We thus judge that this is due to the data segmentation rather than the model assumptions (S2 Appendix) An overview over the correlation values in all setups is given in S1 Table.

**Figure 4.**
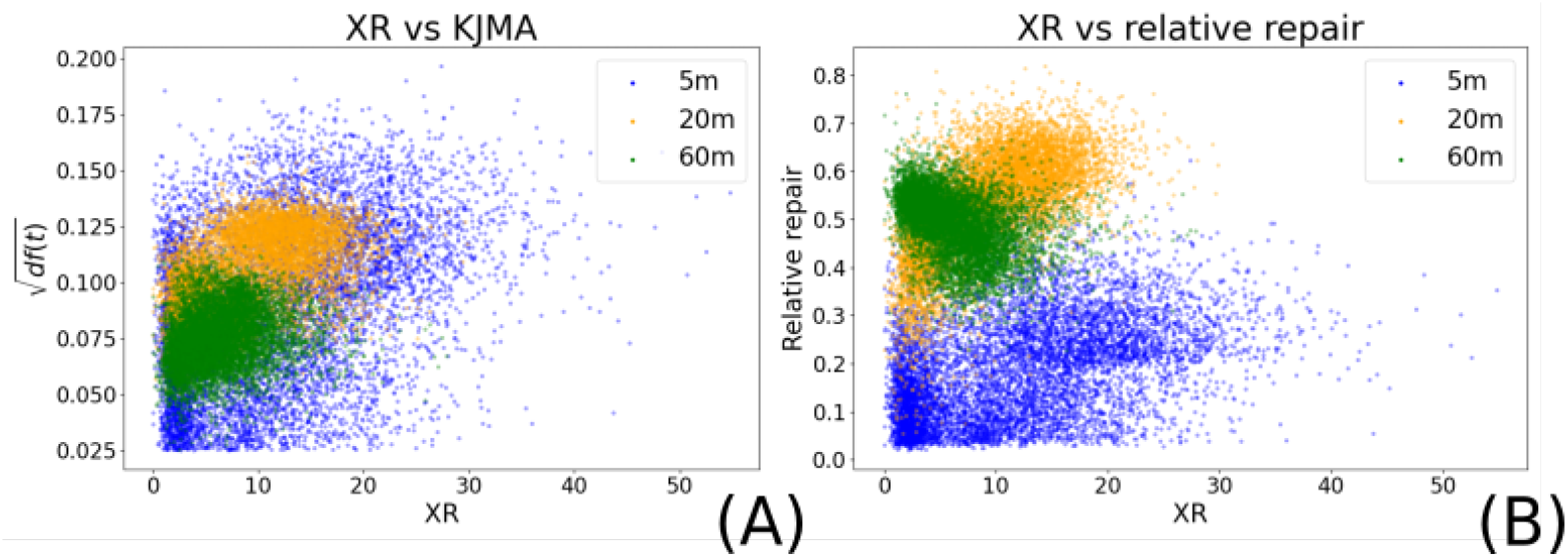
Comparing XR-seq data with model predictions. The values at 5 minutes are given in blue, 20 minutes are coloured yellow, and 60 minutes are green. The plots show that the distance correlation between prediction and XR-seq data is even higher than for the *relative repair rate*. (A) Predicted *relative repair rates* with respect to the XR-seq data in the *TCR* setup exhibit a considerably strong correlation (DC=0.441). Predictions are given as the square root of the model prediction. This reduces the effect of increasing variance with larger derivatives. (B) The *relative repair rate* data as a function of XR-seq values shows a weaker correlation (DC=0.209).

In the way we applied Eq 1, the model assumes that only one mechanism is driving the DNA repair kinetics, i.e. early TCR or late GGR. Using the physical terminology, this results in a constant *nucleation rate*. Whilst this is unproblematic for the NTS and intergenic regions, it is commonly assumed that they are non-exclusive for genes. Interestingly, we can recover the collective effect when averaging the repair evolution for a group of transcripts, e.g. the centre of TCR regions. The beginning of TCR areas is almost solely repaired by a single mechanism, which we assume to be TCR (Fig 5 (A)). The contribution of this pathway is decreasing as a function of distance from the transcription start site (TSS). Although TCR is still involved for lesion removal at the centre and end of the TS, we can observe a strong contribution by a later mechanism (Fig 5 (B) and (C)). We conjecture this to be the effect of GGR. Lastly, lesions in non-TCR regions are only detected by GGR (Fig 5 (D)). This is even the case despite the fact that non-TCR areas also include transcripts. The parameter distribution (i.e. *m, τ*, and *θ*) that create these repair kinetics are exemplified for some setups in Fig 6.

**Figure 5.**
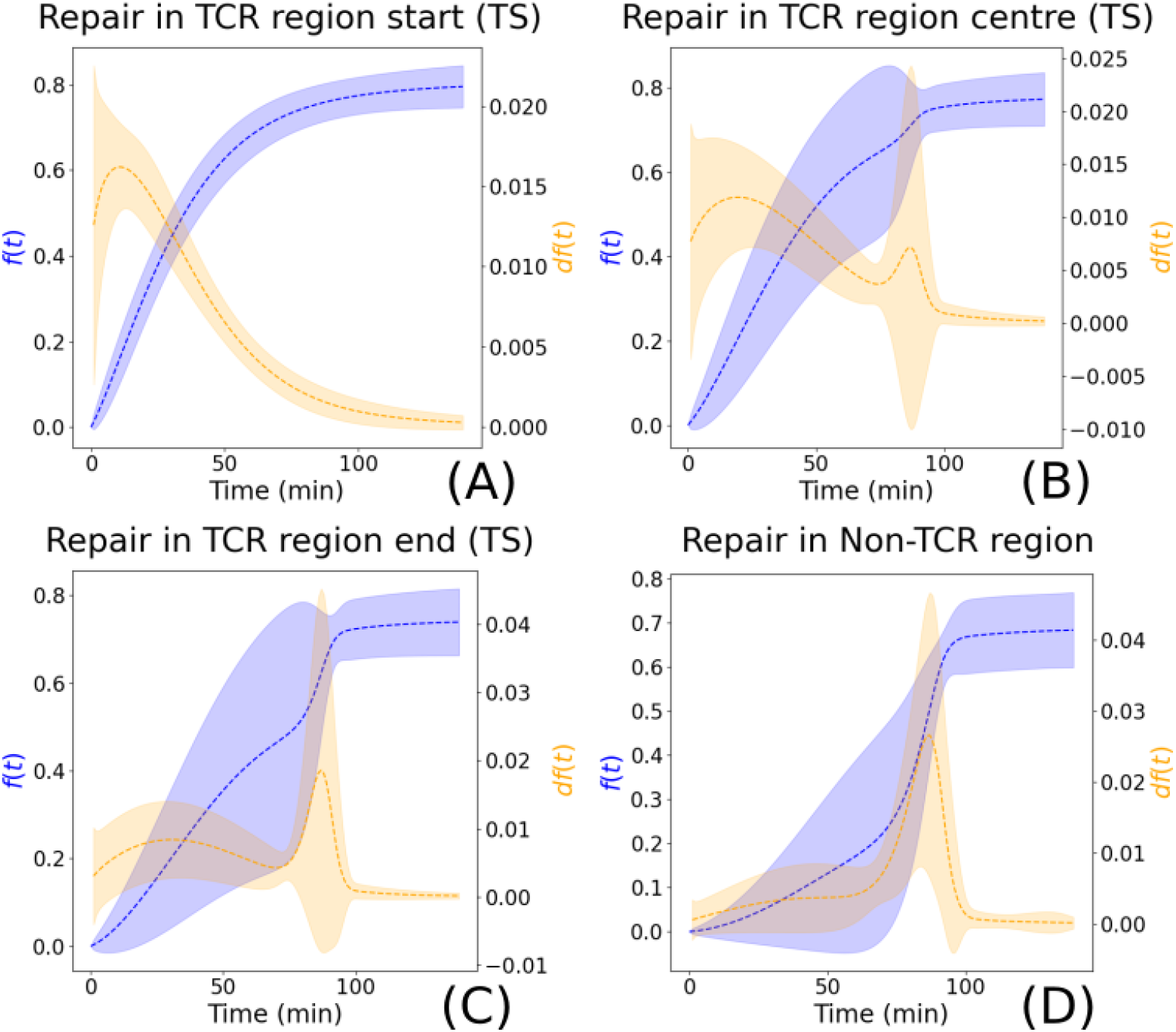
Collective behaviour of genomic regions can recover mutual effect of TCR and GGR. (A) The start of TCR regions is repaired early after irradiation, demonstrating the effect of TCR. (B) At the centre of TCR areas, we can observe the mutual effect of TCR (first peak) and GGR (second peak). (C) GGR’s contribution increases whilst the impact of TCR becomes less important towards the end of the gene. (D) Non-TCR regions are solely repaired by GGR. Therefore, repair is expected at later time points during the process.

**Figure 6.**
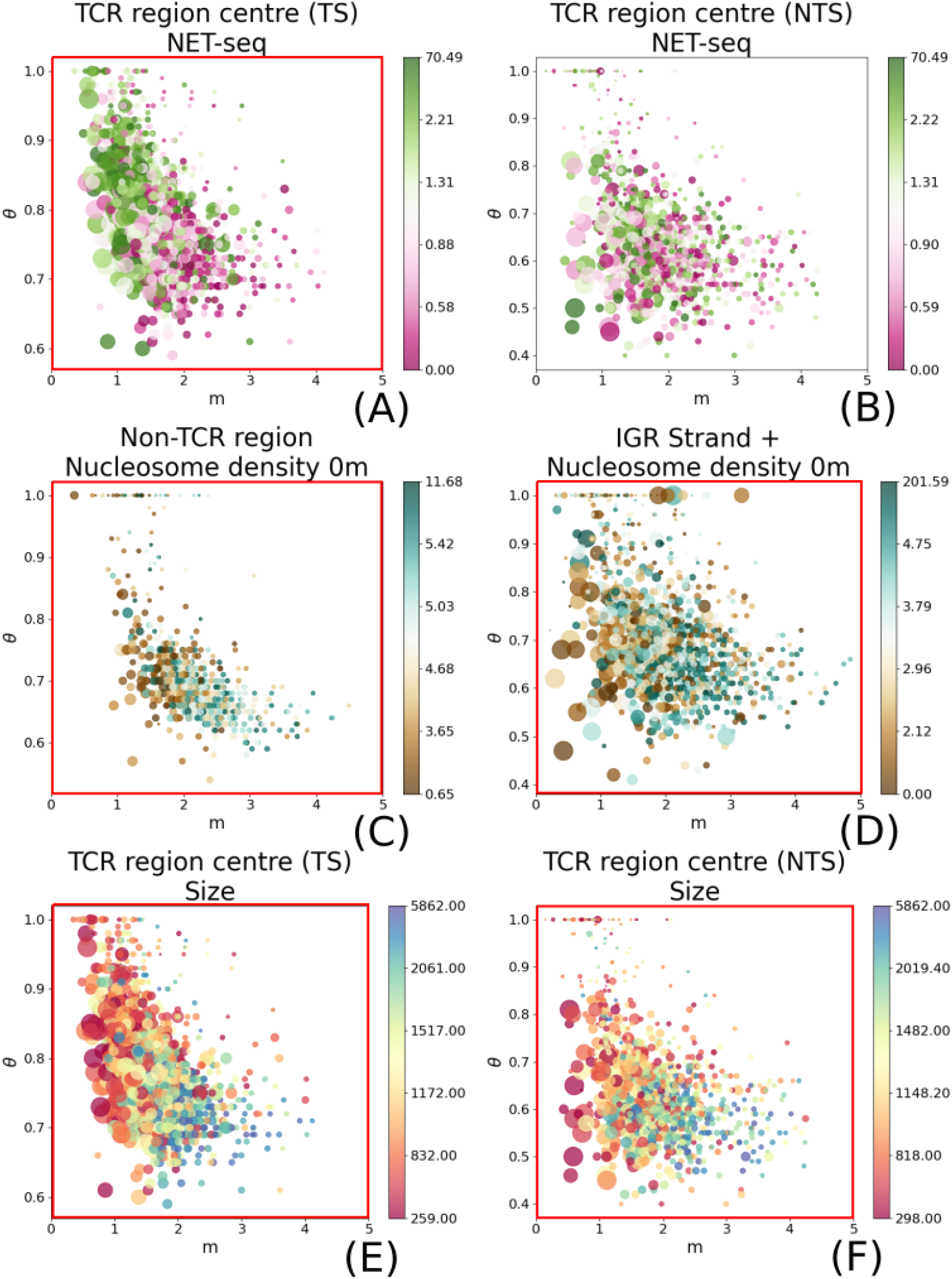
The parameter distribution is coloured with respect to different genomic properties. The x and y-axis give the values of *m* and *θ*, respectively. The size of the circles show 1*/τ* : the larger the circle, the shorter the characteristic time. (A) and (B) are coloured with respect to NET-seq data; (C) and (D) indicate the nucleosome density; (E) and (F) show the distribution with respect to the gene size. Significant correlation is marked with a red frame (*p* value ≤ 0.001 and 90% of the prediction error is below 0.5 in at least three out of five *k*). We define the Watson strand to be +, whereas the Crick strand is −.

Parameter values outside of a reasonable range were excluded from the further downstream analysis. *θ* was set to be between 0.5 and 1 for genic/TCR regions and restricted to 0.4 − 1 for all other; 0.5 *< m* ≤ 6.0; and 20 ≤ *τ* ≤ 200min. It should be pointed out, though, that the number of discarded areas—which had parameters outside the aforementioned ranges—varies depending on the segmentation setup (S2 Table). Motivation and consequences of this approach are explained in S3 Appendix. There are also some regions for which no parameters could be found. However, this concerns less than 10 areas in each experimental configuration. We verified that they did not exhibit any change of repaired lesions. This can happen due to the following reasons. Firstly, the data rectification as defined in Eq 4 sets all data points to the same value. Secondly, the regions did not possess any initial damage. Lastly, they did not possess any pyrimidine dimer sequences where CPDs could be induced (data not shown).

The parameter distributions change considerably over different setups (see examples in Fig 6). Almost all starting areas of TCR regions possess a shape parameter *m* lower than two (figures (A) in S5 Fig to S10 Fig). Moreover, the characteristic time is short. This speaks for efficient repair during early time points, which subsequently flattens out over time (compare with examples in Fig 2). This behaviour is expected since we define TCR regions to be genes that exhibit a strong CPD signal decrease after 20 minutes. It demonstrates that the model parameters are reasonable and match our requirements. Due to the selection of TCR regions, the clear pattern of quick repair (i.e. low *m* and *τ*) vanishes when considering the *gene* setup (figures (H) in S5 Fig and S7 Fig as well as (I) in S6 Fig, S8 Fig, S9 Fig, and S10 Fig). The values are scattered much more broadly over the parameter space.

All other areas show a larger range for the parameter values *m* and *τ*. Non-TCR regions, in particular, have much longer characteristic times than the TS and NTS of TCR areas (Fig 6 (B) and (C)). There is a tendency of 1*/τ* decreasing as a function of *m*; larger circles are more strongly distributed to the left of the figures (low *m*), whilst smaller disks appear predominantly on the right side (large *m*). This is in line with a property of the physical KJMA model which links the turnover rate 1*/τ* to *m*, providing further support for its applicability (see S11 Fig). It is moreover remarkable that the majority of DNA regions can be best described with *θ* ≨ 1; data points are predominantly between 0.6 and 0.9. This could potentially hint to a significant amount of cells that cannot repair their lesions within 120 minutes due to their cell state.

### Correlating Repair Dynamics With Biological Properties

In the previous sections, we showed that Eq 1 describes repair kinetics on a population scale without imposing a particular molecular mechanism. Nevertheless, it can be readily used to investigate the relationship between genomic contexts and the model parameters which explain repair dynamics over time. Hence, we extended the analysis to evaluate influencing factors that indicate relationships *in vivo*.

Previous studies published a large palette of different data types that measure different nuclear properties. We opted for transcription rate, nucleosome density, and gene size as potential influencing candidates. We used the NET-seq signal produced by [16] as a surrogate for transcription rate without UV irradiation. The gene size was measured by [32]. Nucleosome data was acquired after UV treatment by [41].

Correlations with the model parameters were verified with a significance test. We applied a non-parametric two-class *k*-nearest neighbour (*k*NN) approach to allow comparisons between the performance of different *k*NN models. *k* was changed over five iterations. Data was randomly divided into train and test sets, which was repeated 100 times to remove data-specific biases. A correlation was defined to be significant when yielding a p-value (t-test) lower than 0.001% for more than three different *k* in comparison to predictions by a random model. Moreover, it was required that 90% of the models’ prediction error (Eq 12) is lower than 0.5. This corresponds to the expected prediction error of an unbiased coin-flipping experiment.

Correlations considerably differed depending on the genomic area and context (S3 Table). TCR has been repeatedly investigated with respect to transcription rate, and it is surmised to be positively correlated. Hence, higher transcription yields quicker repair, whereas low transcription results in slower lesion removal ([26, 28]). Our model confirms these findings (Fig 6 (A) and (B)). This is true for the *TCR* (beginning, centre and end) as well as the *gene* setup. As expected, the effect was clearly visible on the TS. Surprisingly, though, we find the NET-seq data also weakly correlated with the NTS in the *gene* treatment (see Discussion). It proves that the model can be used to distinguish between distinct regions to provide different levels of detail. Non-TCR and non-transcribed segments seem to be starkly influenced by the nucleosome density (Fig 6 (C) and g(D)). The correlation was clear in all tested setups. There seems to be a weak link to genic areas and the beginning of TCR regions as well. Yet it can be only established for the TS. The clearest results, however, were obtained by gene size. Both, TS (Fig 6 (E)) and NTS (Fig 6 (F)) are clearly influenced. This result was reproduced with all *k*. The gene size is therefore clearly contributing to the lesion removal kinetics. This is an unreported finding for budding yeast to our knowledge. The developed quantitative framework has hence the potential to identify established as well as new interrelationships. The number of *k* values that found a non-random link are given in S3 Table.

We also evaluated more ambiguous links to repair kinetics that have been proposed in the literature. The assessment of Abf1 and H2A.Z histone marker distribution is given in S5 Appendix. We also investigated the impact of the relative distance to the centromere or telomere in order to provide a negative control. We define the relative distance to be the minimum separation to the centromere and telomere divided by the maximum distance possible. Thus, the centre between the two is set to one, whereas the centromere or telomere itself is zero. Indeed, we could not find a significant correlation to any region. The only exception is the TS in the *gene* setup. Yet such a link cannot be found for the *TCR* configuration. This would be expected in such a case since the genes in the *TCR* are a subset of the transcripts in the *gene* setup (S10 Fig).

## Discussion

In this work, we developed a novel computational approach to describe the DNA repair kinetics on a population scale. It is region-specific based on the genome-wide distribution of DNA damages. This required first a new interpretation of the available sequencing data. We introduced a hidden axis that represents autonomous and non-influencing cells. Thus, the one-dimensional signal is now understood as a collapsed two-dimensional grid. By presuming independent repair between and within cells, we can conceptually re-arrange the positions on the grid to create forms. On the basis of this virtual space, we can define ongoing repair as pattern expansion. This permits the application of the KJMA model.

We applied Eq 1 to explain CPD repair kinetics as a function of time. Instead of analysing single time points, we were now able to represent the entire temporal process with three parameters (*m, τ*, and *θ*). This allowed an in-depth comparison of repair dynamics in different areas over the genome. Importantly, it points out that the signal changes non-linearly over time. Whilst this is no new finding by itself—TCR and GGR are commonly seen as acting within different time scales—it has, to our knowledge, not been incorporated in the analysis and prediction of temporal changes in sequencing data. Moreover, the derivative (Eq 3) provides key information about active ongoing repair. It thus permits linking CPD-seq data—showing the DNA damage distribution over the genome—and XR-seq data of excised DNA fragments generated by repair. This provides strong support for the validity of the model. Even though Eq 1 can represent repair only with one mechanism per region, the combined effect of TCR shortly after UV treatment and GGR at later time points can be recovered when considering the average over several areas. The model can be readily used to uncover interrelationships between repair parameters and genomic contexts. The non-parametric *k*NN approach allowed not only finding a significant correlation between model parameters and other biological processes, but also made the results comparable. Our outcomes are consistent with known influencing factors such as transcription rate and nucleosome density. Remarkably, the clearest link was established between the repair dynamics within genes and their length. To our knowledge, this is an unreported finding for budding yeast. In the following sections, we discuss the relevance of our approach and results within the context of previous publications.

### Applying the CPD Repair Model

Several studies proposed temporal models for UV-induced lesion repair on different levels of detail. [27] represented NER kinetics in human cells using a Markov-Chain Monte-Carlo approach. It explains the removal of 6-4 photoproducts on a single-cell scale through the random and reversible assembly of repair complexes. A similar model was proposed by [34]. Interestingly, though, they derive very different conclusions, as they suggested that random or pre-assembly of repair proteins is unfavourable. Despite a great level of detail of both models, they are incapable to make region specific predictions. Moreover, as both models are based on microscopy data, they do not explain temporal changes in genome-wide sequencing data on a population scale. A Monte Carlo approach to explain the damage distribution and subsequent repair induced by ionised irradiation in a single cell was proposed by [38]. It also incorporates the collective effect of NER and base excision repair (BER), therefore accounting for different and potentially competing mechanisms. It should be stressed that the lesion type is considerably different. Again, the predictions are location unspecific. [40] provided a different angle by presenting a protein-protein interaction landscape of NER components in yeast. Predictions about the repair efficiency in different regions were not established. To our knowledge, our model is the first that accounts for region specific changes in population-based data.

In order to apply Eq 1 and to re-group repair regions to patterns, we presumed independent repair kinetics between and within cells. It is not far-fetched to conjecture that each yeast cell repairs DNA damages autonomously. *Saccharomyces cerivisiae* are single-cell organisms and should thus react independently to DNA damage. Moreover, yeast cultures were grown in rich medium after UV treatment, precluding any limitations for growth ([28]).

Our second presumption surmises independent repair dynamics within a cell. This is based on two observations. Firstly, we assume that the spatial effect of lesion removal kinetics decreases as a function of distance. Hence, the farther away the CPD positions, the smaller the impact on each other. This is justified by the relatively small area of lesion removal (≈ 30 nt ([7])) and the notion of chromatin interaction domains (CIDs) ([17]). Secondly, it has been reported for *Caenorhabditis elegans* that a UVC treatment of 100J/m^2^ induces 0.4 to 0.5 CPDs per 10kb ([31]). A similar UV dose (125 J/m^2^) was used by [28]. Taking this as a reference, we presume that a comparable dose of UVC induces a corresponding number of CPDs in budding yeast. It is thus unreasonable to expect more than one lesion per CID per cell, as they are commonly less than 10kb in *Saccharomyces cerevisiae* ([17]). It is true, though, that this could be species-dependent. Due to the lack of other studies, we take it as given that lesion removal does not affect each other within a single cell.

The independence assumptions permitted the application of the KJMA model. As we surmise that the sequencing data contains a hidden axis, we represent repair on a grid. Thus, it would be expected that the found shape parameter indicates a two-dimensional space, i.e. *m* − 1 = 2. However, as reported above, *m* exhibits a large range. We do not presume that such a deviation is only caused by noise. Instead, a similar behaviour can be observed when allowing the growth speed *G* to be larger during earlier time points rather than later in the process, and vice versa. Implicitly, this incorporates that *G*(*t*) is non-constant in time. *m* can be interpreted to speak for the time-dependence of the process instead of a particular dimension ([8, 35]). Low values represent quicker repair in the beginning rather than in the end. A large *m* indicates that *G* increases later on. Eq 6 explains the dependence of the observed expansion speed on *G*(*t*) and *m*.

The nucleation rate *n*, though, is presumed to be constant. As explained above, this has as a consequence that the framework models repair with only one mechanism per region. There is a scientific consensus that intergenic regions and the NTS can be only repaired by GGR. For the TS of genes it is nevertheless surmised that TCR and GGR can act collectively. On a population scale, this would likely result in the repair rate to contain two peaks over time. TCR would be observable in the beginning and subsequently abate. GGR is supposedly acting later during the process. The collective effect of TCR and GGR in a genic area can be recovered by taking the average over an entire group, e.g. the beginning of TCR regions. Despite assuming similar kinetics, we presume that the noise in the process should lead to a representation by either TCR or GGR in a ratio comparable to their respective repair contribution. It should be highlighted that Eq 1 can be easily adapted to represent heterogeneous repair by defining *n*(*t*) as a function of time. However, any parameter estimation of such a function would be merely based on guesses due to the sparse temporal data resolution. We followed the principle of Occam’s razor and opted for a simpler model. The production of CPD data with smaller time steps could permit such an estimation.

### Correlating CPD Repair Dynamics With Genomic Properties

The parameters of Eq 1 can be used to set repair into the context of other biological and nuclear properties in the cell. As databases provide a large variety of data signals, it is reasonable to opt for a data driven approach. However, finding such a correlation is not straightforward. Many of the NGS histogram distributions peak sharply around a low value whilst also including far distant outliers. Moreover, it can be assumed that sequencing signals include a large amount of noise. This makes it difficult to find a correlation using a continuous regression approach. We circumvented these issues by transforming the mapping into a binary classification problem. This reduces the impact of noise. Due to using equally many values for both classes, we removed any distribution specific bias.

Another requirement was comparability between results. It is a known fact that the performance of machine learning models can vary strongly depending on the number of parameters or used architecture ([10]). Through using the non-parametric *k*NN approach, we could provide equivalent treatment for all setups. We also mitigated the impact of *k* by applying different values in a reasonable range, i.e. *k* ∈ {5, 10, 20, 50, 100}. Thus, we did not rely on any particular parameter setting. We are aware that parametric models can be efficiently implemented for a variety of tasks, as it has been recently shown with Alpha Fold 2 ([21]). Moreover, architectural biases of parametric models could be possibly reduced by systematic parameter searches. Nevertheless, it should be emphasised that this study did not intend to find the best performing machine learning model. Rather, we aimed to show non-random correlations and therefore indicate potential repair influences. Some researchers even conjecture that non-parametric models could be generally better performing ([33]). We conclude that the *k*NN approach is a sensible choice.

We want to provide some further intuition for the functioning of the *k*NN method (see Methods and Materials) with the example of the transcription length. Fig 7 shows the learnt function and the prediction error distribution for the correct and random model, respectively. The correct mapping finds a distribution pattern with big genes predominantly distributed to the centre right, whereas small genes are found in its periphery (Fig 7 (A)). As expected, this pattern is destroyed in the random mapping (Fig 7 (C)). The error distribution for the true *k*NN is equally large for values that are over and underestimated (right histogram in Fig 7 (B)). This speaks for an unbiased mapping. We also want to point attention towards the spatial distribution of the prediction error. Red circles mark data values which were incorrectly classified as large genes, whereas the blue points are parameters that were wrongly associated with a small size. The distribution of the red and blue disks follows our expectations from the learnt map. The wrong classification could indicate noise in the data set or unknown information that cannot be represented. Compared with Fig 7 (D), it is clear that these trends vanish in the random model. However, the prediction error is surprisingly low (0.4). A repetition over 100 iterations is therefore indispensable to find significant links and to remove data specific biases.

**Figure 7.**
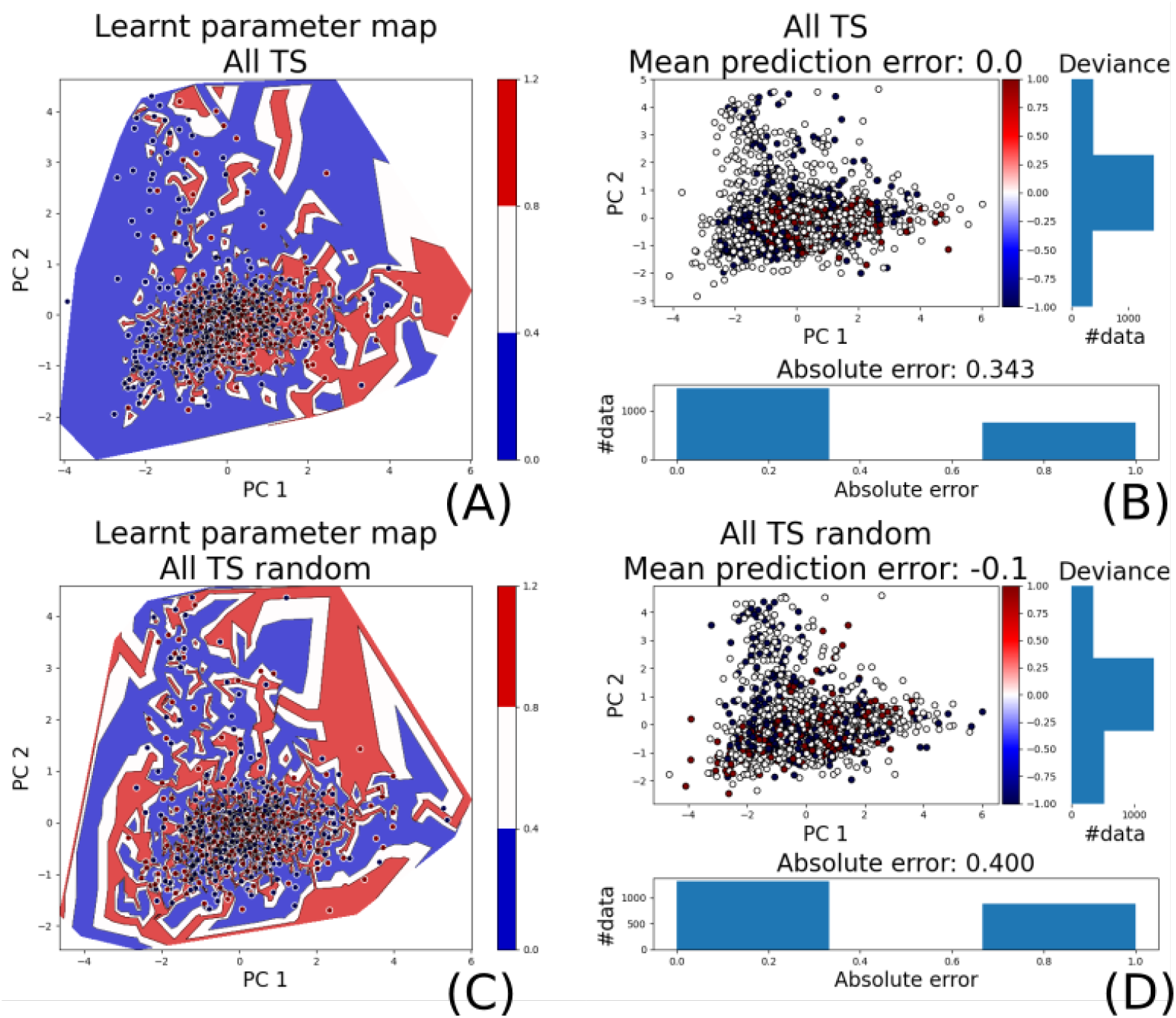
Example of the learnt function between model parameters and genomic context. (A) The learnt parameter distribution and the associated class for the true model after applying a principle component transformation. The x and y-axis give the first and second principle component, respectively. Red represent large genes, whereas blue shows low values. (B) The error distribution for the predictions follows the expected outline given by the learnt function in (A). The blue and red circles give values that were classified as short but were actually large and vice versa, respectively. White points are correctly classified. The right bar shows the error distribution along the colour axis, i.e. over estimated, correctly classified, and underestimated values from top to bottom. The lower histogram shows the distribution of overall correctly and incorrectly classified values. (C, D) The parameter learnt parameter map and the error distribution of the random model.

### Analysing Genomic Properties Which Influence Repair Kinetics

TCR has been identified as a rapid repair pathway on the TS. Intergenic regions and the NTS exhibit significantly slower lesion removal, which was demonstrated on the genomic scale in yeast and human cells ([18, 19, 26, 28, 29]). It remains an unsolved quest to find an interrelationship between TCR efficiency and transcription rate. Whilst the two parameters are indeed assumed to be correlated ([26, 28]), some studies point out that TCR repairs CPDs efficiently at nearly all genes including those with a low transcription ([29]). An in-depth analysis is still missing, and there is no clear consensus on how transcription rate is affecting repair. In this work, we compared the model predictions to gene expression. Our analysis clearly shows a significant correlation on the TS and is therefore supporting the common assumption (S5 Fig).

As repair proteins need to recognise and repair lesions on the DNA, it is conjectured that chromatin organisation can significantly modulate the efficiency of CPD repair ([28, 29, 41]). However, previous studies were mostly scrutinising the positioning of damage at nucleosomes. CPD removal was shown to be less efficient at the dyad of strongly positioned nucleosomes in yeast ([28]). Moreover, GGR on the NTS was asymmetrically inhibited in yeast and human cells with respect to the position within the nucleosome([29]). Even though nucleosome occupancy after UV treatment was already previously probed, the potential relationship of these data with CPD repair was not directly addressed ([41]). Our results demonstrate a significant correlation between repair and nucleosome density in intergenic and non-TCR regions (Fig 6 (C) and (D)). We also discovered a clear influence on the beginning of TCR areas (S6 Fig).

Unexpectedly, our outcomes show a strong correlation between transcription length and repair. Differences in transcription shutdown and restart after UV treatment relative to gene size were previously reported for human cells ([42]). Both transcription regulation and efficient repair are necessary to orchestrate an effective cellular response to UV light. The restart of transcription to pre-irradiation levels is an important step at the final stages. However, a direct evaluation of lesion removal with respect to gene size was not performed. To our knowledge, this is a new finding for CPD repair in yeast. Due to our data pre-processing (Eq 4), we can rule out that the result derives only from the fact that larger areas have a greater potential to include more damage. This is true due to two reasons. Firstly, we normalised the CPD value in each bin (e.g. beginning of the TS) by the number of pyrimidine dimers in the sequence as described in [28]. Secondly, and more importantly, we want to point out that the quotient in Eq 4 lets any length dependence and normalisation of the binned data vanish. Therefore, the values become automatically comparable due to the design of Eq 4. It should be mentioned, though, that the regions of interest can become rather small when segmenting the gene into subareas. Influence or noise from neighbouring areas can not be excluded. However, due to the fact that the same result can be obtained in the *gene* setup (Fig S7 Fig), we presume that it represents a genuine feature of the CPD removal mechanisms in yeast cells.

Lastly, we investigated a potential link to the distance relative to the centromere and telomere depending on which was closer to the region of interest (S10 Fig). We could not find a non-random link between the parameters and the position. The only exception was the TS in the *gene* setup. However, the error distribution was very close to the random distribution (data not shown), and the result could not be reproduced with the *TCR* configuration. This would be expected as the TCR regions are a subset of all genes. We therefore do not assess it as an important result.

In conclusion, our work opens interesting perspectives for future research on DNA repair mechanisms and influencing genomic factors. New experimental data with increased temporal resolution will help to refine the model and analysis. The approach can be similarly used for other organisms including human cells. Moreover, it can be readily applied to sequencing data of any nuclear process that can be represented as a two-state system, and it is not restricted to repair.

## Methods and Materials

### Data Treatment

All experimental data that were analysed in this study comes from public databases (see overview in Table 1). CPD-seq data was taken from [28]. It contains two time courses with samples taken at *t*_1_ ∈ {0, 60} min and *t*_2_ ∈ {0, 20, 120} min, respectively. The location of transcribed areas was taken from [32]. Data signals were partitioned into different segments, depending on the experimental procedure (see schema in Fig 3). We distinguished between two schemes. The *gene* setup categorises the data into TS and NTS as well as the plus and minus strand of intergenic regions. The *TCR* configuration uses the notion of TCR regions (see S1 Appendix). All areas that are non-TCR regions are grouped together. We furthermore partition the TS and NTS of TCR regions into beginning, centre, and end. For the non-TCR areas, we combined both strands to one group. Hence, for finding the parameters of Eq 1 of non-TCR regions, we used two data points per time point (each representing repair on one strand).

**Table 1.** Overview over the data sets that were used in this study.

| Property | Strain | Data type | UV Dose | Reference |
| --- | --- | --- | --- | --- |
| CPD | BY4741 (WT) | CPD-seq | 125 J/m<sup>2</sup> | [28] |
| CPD repair | Y452 (WT w.r.t. repair) | XR-seq | 120 J/m<sup>2</sup> | [26] |
| Abf1 | BY4742 (WT) | ChIP-seq | 100 J/m<sup>2</sup> (0min) | [41] |
| H2A.Z | BY4742 (WT) | ChIP-seq | 100 J/m<sup>2</sup> (0min) | [41] |
| Nucleosome distr. | BY4742 (WT) | MNase-seq | 100 J/m<sup>2</sup> (0min) | [41] |
| Transcription rate | YSC001 | NET-seq | - | [16] |

CPD-seq fragments were normalised by the number of available pyrimidine dimers, as explained in the supplementary material of [29]. The damage distribution was subsequently transformed into relative repair in area *a* through

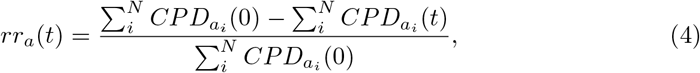

where *N* denotes the size of *a*, 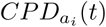 is the normalised CPD signal at time *t* and locus *i* in area *a*, and *t* ∈ {20, 60, 120}min. *a* is any of the previously described regions (e.g. the start of the NTS) depending on the experimental configuration. We additionally take it for granted that no new CPD lesions can be induced during repair. Hence, all data points were enforced to be greater than or equal to zero and monotonously increasing as a function of time. This was defined by

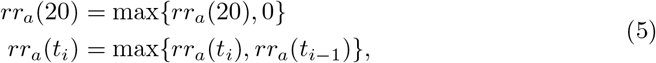

where *t*_*i*_ *∈* **t** = (20, 60, 120).

To investigate the link between the repair evolution and other genomic contexts, we included a broad and heterogeneous data set in our analysis. Several studies proposed a variety of influencing factors; among others, the correlation of TCR with transcription rate ([26, 28]), as well as the influence of nucleosome positioning, histone marks, and other repair related proteins ([41, 43])(Table 1). All sequencing data were averaged over the size of the investigated area. In setups during which we scrutinised different subregions of genes (i.e. start, centre, and end), all of them were linked to the same value to smooth out the potential influence of noise. For example, the beginning, centre, and end of a TCR region were all associated to the same transcription rate. Moreover, both strands were compared to the same data, e.g. the TS and the NTS were related to the same nucleosome density. We noticed that the NET-seq signal amplitude decreases as a function of distance from the TSS (S12 Fig). This could possibly induce a gene size-specific bias that is not removed by taking the average over the transcription length. We could verify, however, that the NET-seq data strongly correlates with independently probed Pol2 ChIP-seq data ([13])(S13 Fig, S14 Fig). We therefore assume that it reasonably represents transcription rate, whilst allowing a direct comparison to the results obtained by [26] (see also S4 Appendix).

With the exception of nucleosome density, all biological data values are seemingly Poisson distributed. Some of them strongly peak around a low value, but contain large positive tails. To remove a potential bias introduced by outliers, we limited our analysis to the lower 95th percentile. As this procedure was applied to all data (except nucleosome density), we did not introduce a bias towards a certain model. Rather, we improved comparability. The only exception is the MNase-seq signal, as it is approximately normally distributed. We consider that trimming could introduce a bias rather than removing one.

We furthermore assessed the biological data distribution for different region models (e.g. the beginning of the TS and NTS of TCR regions) after removing parameters outside defined ranges as well as excluding the outliers. We could verify that in most setups, all histograms were similarly shaped, and did not contain any apparent bias towards a genomic area. However, the data distribution of Abf1 and H2A.Z data was different in intergenic regions compared to transcribed areas. This is further evaluated in S5 Appendix.

### The KJMA Model

The KJMA model describes the phase shift (*α* to *β* phase) of solids at constant temperature. This is expressed in the fraction of successfully transformed *β* particles with respect to the total volume. In this section, we want to briefly introduce the physical model. All terminology refers to its physical context, and it does not express biological properties. For a comprehensive analysis, see [2–4, 12, 20, 24, 37].

It is assumed that atoms jump or migrate from an *α* matrix to a *β* particle through random nucleation. The new *β* particles grow at a certain speed *G* and stop when they impinge on each other. Strictly seen, this permits the phase shift only to occur in so far untransformed material. The mathematical treatment is simplified when instead considering the extended volume first. This means the fraction of transformed volume is computed as if the entire sample were still in phase *α*. Let *f* (*t*) ∈ [0, 1] be the real volume fraction of transformed *β* particles. Moreover, *V*_*β*_ and 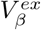 are the real and extended volume, respectively. 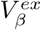 is computed based on the number of transformed nuclei that expand as a volume *v*. We consider the time interval 0 *< η < t*. New nuclei are created with a rate *n*. Hence, the number of new nuclei *N* in a given volume *V* that are created within *dη* are described by *N* = *V ndη*. When we assume *G* to be constant, the transformed volume by a single particle within a time period *t* − *η* can be described by

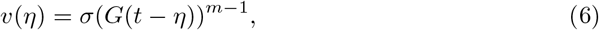

where *m* is the Avrami exponent which determines the dimension in which growth is modeled (e.g. *m* − 1 = 3 for three-dimensional expansion). Furthermore, *σ* describes the shape, e.g. *σ* = 4*π/*3 for spheres. Taking it all together, we write

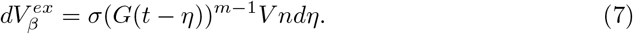

Integrating both sides yields

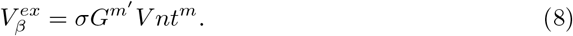

However, as mentioned before, the transformation can actually only happen in untransformed material. Eq 7 can be updated to

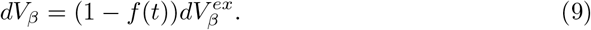

When re-arranging and integrating Eq 9, we obtain

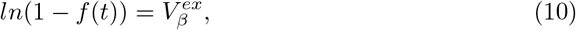

which can be brought into the general form of the KJMA model

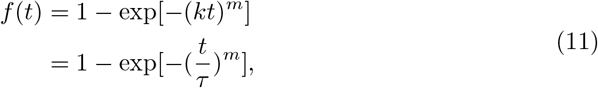

with *τ* denoting the characteristic time.

### Parameter Optimisation and Definition of Correlation

We presume that temporal changes in CPD-seq data can be explained by Eq 1. Parameters *m* and *τ* were determined by applying Eq 2. During the parameter search for *θ*, we started with 0.5 as minimal value for transcribed/TCR regions and 0.4 for all other. We increased it thereafter by Δ*θ* = 0.01, until we reached *θ* = 1.0. We hence constrained the parameter search such that at least 40% of all cells—and in transcribed/TCR regions at least 50%—must be able to repair their lesions. We conjecture that data within this range is reasonably well described by Eq 1.

We pursued a data driven approach in which we estimated a function between the parameters of Eq 1—namely the shape *m*, the characteristic time *τ*, and the maximum repair fraction *θ*—and other biological contexts. For the actual implementation, we used the conversion rate 1*/τ* = *k*. We grouped biological data into high (class *c* = 1) and low values (*c* = 0), such that both classes contained the same number of samples. To train the machine learning model, the input values **x** = (*m, k, θ*) where normalised such that every dimension was normally distributed with zero-mean and a standard deviation of one. We opted for a *k*NN approach with *k* ∈ {5, 10, 20, 50, 100}. It compares an unknown input 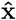 to the *k* closest values of a known data set {**X, c}** to predict class *ĉ*. We opted for the Euclidean distance as similarity measurement. Here, the *i*-th row of **X** is **x**_**i**_ = (*m*_*i*_, *k*_*i*_, *θ*_*i*_), and *c*_*i*_ is the associated class in **c**. *ĉ* is determined by a majority vote. For example, if more than 50% of the *k* neighbouring values are classified as group *c* = 1, then *ĉ* is predicted to be group 1 as well. *k*NN is categorised as non-parametric model which permits the comparison of different results. The model performance was measured through calculating the prediction error

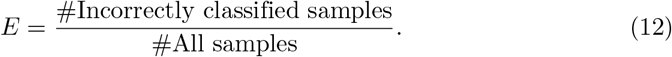

This was compared to a random baseline model, for which the classes were randomly shuffled to a given parameter triple (*m, k, θ*) during training. Data was arbitrarily partitioned into learning and testing data sets. Every experiment was independently repeated 100 times to reduce the effect of any potential bias. We consider an interrelationship to be important if the prediction error is significantly lower than the accuracy of the random model (*p <* 0.001% of a one-sided t-test). Moreover, we require that 90% of the prediction errors are below *E <* 0.5 which is the expected outcome of an unbiased coin-flipping experiment. This significance must be found in three out of the five evaluated *k* to indicate an interrelationship.

## Acknowledgments

LZ was supported by the CEA NUMERICS program, which has received funding from the European Union’s Horizon 2020 research and innovation program under the Marie Sklodowska-Curie grant agreement No 800945. The work was supported by the Fondation ARC (PGA1 RF20170205342). We thank Zoë Slattery for proofreading the manuscript.

## S1 Appendix

### Determine TCR Regions

In this study, we distinguish between two genomic segmentation schemes: the *gene* configuration and the *TCR* setup. The former is constructed as follows. Coordinates for transcribed regions were taken from [32]. This is set to be the TS. The area opposite of the TS is the NTS. All other segments are defined to be intergenic or non-transcribing. Here, we distinguish between Watson (positive) and Crick strand (negative). This permits us to show that there is no strand-specific bias. For the *TCR* setup, we introduce the notion of TCR regions. They are defined to be transcripts that repair more or equal to 20% of their initial repair within 20 minutes. Approximately 89% of intergenic regions possess repair rates lower than 20% during the same time span (S1 Fig). Hence, we can have an increased confidence that genic regions with quicker repair are supported by TCR, leading to a more than 1%-point decrease of CPDs per minute. For all other genes, we cannot exclude that they are only repaired by GGR, as it can function genome-wide. It is important to highlight that the observed CPD decrease is not uniform along the transcript. The end of the TS is seemingly less efficiently repaired. We require that only the first third of the gene after the TSS must possess more than a 20% decrease of damage within 20 minutes.

## S2 Appendix

### Finding a Relationship Between XR-seq Data and Repair Rate

As the derivative of Eq 1 represents the *relative repair rate* at a given time, we conjectured that it should correlate with XR-seq data. This is due to the fact that the signal shows the distribution of excised nucleotide sequences that were produced during DNA cleavage. It is surmised that they are quickly degraded, i.e. within a five minutes ([26]). However, this interrelationship is not necessarily linear, which speaks against the usage of Pearson’s correlation. The distance correlation (DC) comes as a remedy by correlating the distance of data points in a set to each other rather than the data itself. It ranges from zero to one, with zero indicating independence, whereas one indicates that the linear subspace between the data sets is equal. It is calculated as follows. Distance matrices *A* and *B* contain all pairwise distances, i.e. *{A}*_*ij*_ = ||**a**_**i**_ − **a**_**j**_||_2_ and *{B}*_*ij*_ = ||**b**_**i**_ − **b**_**j**_||_2_. Here, **a**_**x**_ and **b**_**x**_ (*x* ∈ *{i, j}*) denote data points in sets *𝒜* and *ℬ*, respectively. ||…||_2_ is the Euclidean distance. Each set contains *n* data points. Subsequently, *A* and *B* are double-centred. With the definition of the sample distance covariance

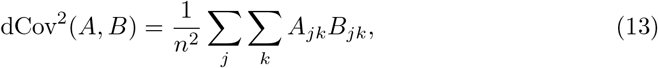

as well as the sample distance variance

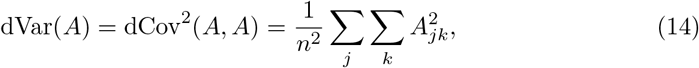

, we can introduce the DC:

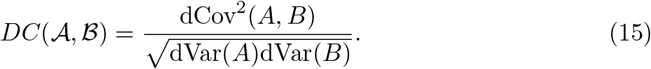

The DCs for the different setups are given in S1 Table. In order to compare the values of Eq 3 to the CPD data, we transformed first the signal with respect to Eq 4. This yields relative repair at three time points, i.e. 20, 60 and 120 minutes. The *relative repair rate* is determined by

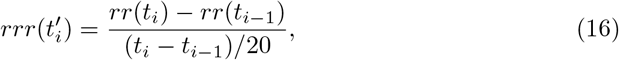

where *rr*(*t*) denotes relative repair (Eq 4). *t*_*i*_ ∈ **t** = (20, 60, 120) and 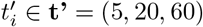. The values must be re-scaled to the same time step to make them comparable. However, the CPD decrease within the first 20 minutes is relatively small for most areas. Finding a *relative repair rate* per minute results in an almost flat line. All values represent therefore repair within 20 minutes. This makes the time points comparable whilst avoiding having too small values for *rrr*(5).

Despite the fact that we determined the DC for all time points—i.e. 5, 20, and 60 minutes after repair—it is intuitive to see that *relative repair rates* are more heterogeneous when taking all time points together. We therefore consider only the DC of the entire data set. Astonishingly, the model predictions of the *TCR* setup are better correlated with the XR-seq signal than the *relative repair rate* data. It should be mentioned, though, that the *gene* setup yields a substantially lower correlation. Since this is true for both, the prediction and the data, we conjecture that this is merely caused by the data partitioning.

## S3 Appendix

### Data Transformation and Selection

In order to make our results comparable, we followed the signal analysis described by [29]. However, we used three bins (beginning, centre, and end) instead of six. We also converted the data such that it represents repair instead of damage. This allowed a straightforward application of Eq 1. We additionally required that repair is greater than or equal to zero and monotonously increasing as a function of time. Some studies propose the notion of *dark* or *delayed* CPDs in human cells, which occur after UV treatment ([11, 45]). However, to our knowledge there has not been a consensus over how *delayed* CPDs occur and influence repair dynamics. As we assume the biological process as well as the data probing itself to induce a considerable amount of noise, we prefer the interpretation that these data points should be rectified, rather than they represent damage created after irradiation.

In order to find potential groups that show similar repair dynamics, we compared the distribution of the model parameters against each other. Depending on the chosen segmentation and the type of genomic region, we found two to four clusters which were predominantly determined by the shape parameter *m*. When investigating the repair dynamics in detail, we found that all groups except one produced a switch-like behaviour (S2 Fig(B)). Whilst this could be a genuine property, we conjecture that this comes from the applied data normalisation. As discussed before, we require that no new lesions can be induced after irradiation. However, almost exclusively all regions with *m >* 6.0 originally possessed larger CPD signals after 20 minutes than directly after irradiation. During the data transformation, this data point was hence set to zero. Due to the form of Eq 1, the lesion removal mechanisms is seemingly acting exclusively between 20 and 60 minutes. As we gauge that this results solely from the noise in the data, we subsequently excluded these regions. In some cases, we could also find a grouping which was driven by the characteristic time *τ*. Large values only occurred in on the NTS or non-TCR/non-transcribed regions. We observed that these areas were all comparatively small, i.e. less than 300 base pairs (bp). Therefore, they are very susceptible to noise and processes from neighbouring regions. Instead of requiring a minimal length, we limited the range of *τ* to keep as many areas with potential useful information as possible. We assumed 200 minutes to be a sufficient time range for CPD repair to occur in yeast. All parameter ranges were set as follows: *m* ∈ [0.5, 6.0]; *τ* ∈ [20, 200]; and *θ* ∈ [0.5, 1.0] for genes (TCR regions) and *θ* ∈ [0.4, 1.0] in non-transcribed (non-TCR) areas. The number of remaining regions that fulfilled the set requirements changed considerably depending on the experimental setup. An overview is given in S2 Table. The TS was in almost all cases included in the subsequent analysis, although the end was more often outside the defined parameter ranges than the beginning and centre. Surprisingly, only around half of the NTSs met the requirements (both setups). The numbers are even worse for intergenic regions (both setups). Here, approximately a third of all non-transcribed/non-TCR areas were considered in the downstream computations. We hypothesise that NTS and intergenic regions exhibited frequently later repair times. This has been also noted by [26] and [28]. Thus, we conjecture that the noise in the system can likely appear CPD levels at 20 minutes to be higher than after irradiation. Considering additionally the few time points, the data fitting of Eq 1 predicts no repair until 20 to 60 minutes, whereas all CPD decrease appears exclusively afterwards. This results in the aforementioned step-like behaviour. As discussed before, we assume this to be rather unlikely. Analysing the repair dynamics in these regions could provide additional information. It cannot be excluded that repair occurs in a short and well defined time frame, e.g. only between 20 and 60 minutes. This could be possibly driven by the accessibility for involved proteins or similar. We hope that future research is inspired to repeat this analysis with a CPD-seq time course that has a finer temporal resolution.

## S4 Appendix

### Comparison of Transcription Rate and Gene Size

It is usually assumed that the size of genes does not influence the frequency with which they are transcribed. Thus, both parameters are expected to be independent. During our analysis, we noticed that the NET-seq signal amplitude decreases as function of distance from the TSS (S12 Fig). This could possibly induce a size-specific bias if the decline occurs repeatedly within a specific distance, e.g. 500 bp from the TSS. In this case, it affects smaller genes more strongly than large genes. In order to gauge the bias’ impact, we compared the NET-seq signal with Pol2 ChIP-seq data ([13]), which we assume to represent transcription rate to a reasonable degree. We can verify that Pol2 ChIP-seq does not exhibit the same declining trend after the TSS. Indeed, NET-seq shows a larger correlation with respect to gene size (DC=0.321) than Pol2 occupancy (DC=0.224, S13 Fig). However, we can establish a rather strong interrelationship between NET-seq and Pol2 ChIP-seq data (DC=0.75, S14 Fig). When we scrutinised the link to repair, we divided all genes into two groups with high or low transcription rate. We can verify that the majority of is still within the same group, independent of the use of NET-seq or Pol2 ChIP-seq data. Therefore, the conclusions about the relationship between transcription and repair remain nevertheless sensible. We opted to use NET-seq data to permit a direct comparison with the results from [26].

## S5 Appendix

### Analysing Repair Kinetics in Context of Abf1 and H2A.Z Distribution

Different chromatin features including transcription factor binding sites or histone variants and modifications can affect CPD repair. An essential role in GGR recognition is allocated to the Rad7-Rad16-Abf1 protein complex. Yeast strains with respective gene deletions are incapable to repair lesions in non-transcribed regions and are inviable under genotoxic stress ([7, 46, 47]). Abf1 binding was also proposed to inhibit CPD formation ([28]) and to influence GGR kinetics ([46, 47]). Moreover, TCR and GGR are both reportedly influenced by multiple histone modifications and variants ([1]). Binding sites for the hypothesised GGR-complex are flanked by H2A.Z histone variant-containing nucleosomes ([41]). However, a direct relationship between lesion removal and Abf1 occupancy or H2A.Z distribution has not been investigated. Building up on previous work, we presumed particularly strong correlation in intergenic regions. Abf1 and H2A.Z distribution was probed after UV treatment by [41].

The experiments with Abf1 yielded a mixed bag of results (S8 Fig). When considering the *TCR* setup, we found a non-random correlation with the repair dynamics in intergenic regions for all *k*. Only when including a larger number of neighbours (*k* ≥ 20), the centre and the end of TCR regions showed a significant deviance to the random model. This behaviour was inverted in the *gene* configuration. The parameters of the TS and NTS could be linked to Abf1 occupancy for all *k*. For *k* ≥ 10, the negative strand of intergenic areas was also clearly showing a non-random correlation. However, we could not find a significant influence for the positive strand of non-transcribed regions before *k* = 100. This is surprising, as we would expect both strands to behave similarly. Whilst this could hint to a strand-specific bias, it is likely that the influence of Abf1 at intergenic regions in the *gene* setup is weaker than in the *TCR* configuration. A correct definition of genomic groups is hence clearly important to put the results into the right context. Abf1’s role is associated with GGR ([41, 46, 47]) as well as transcription regulation and replication ([23, 44]). It is intuitive that due to its multifunctional involvement it is indirectly affecting a broad variety of regions.

The outcomes for H2A.Z were similarly ambiguous (S9 Fig). We found a clear link at the TS of TCR regions and genes. The correlation was significant for all *k*. In the *TCR* configuration, this was accompanied by a non-random link for the NTS with *k* ≥ 10; and with larger *k* also in non-TCR regions (*k* ≥ 20). On the other hand, we were unable to find a significant correlation for both strands in intergenic areas for the *gene* setup. Nonetheless, there was a definite interrelationship for all *k* between the histone marker distribution and the repair dynamics at the TS and NTS. This indicates a non-negligible role for H2A.Z modification during lesion removal at active genes. This is unsurprising giving its regulatory role in gene expression ([14]). However, a correlation with GGR might be less strong.

To put the results into context, it is important to mention that the distribution of Abf1 and H2A.Z data was different in intergenic regions compared to transcribed areas. This was especially visible in the *gene* setup. It has been previously reported that Abf1 binding sites tend to colocalise with CID boundaries, which are usually found in intergenic regions. In the same paper, it was also proposed that they are flanked by H2A.Z-containing barrier nucleosomes ([41]). Assuming that Abf1 is necessary for GGR ([7, 46])—and therefore plays a specifically crucial role in intergenic regions—it is not surprising that the histogram exhibits different distributions for genes and non-transcribed areas. Since the distributions are still similar between strands, we do not consider this as having a strong influence on the final conclusions.

## S6 Appendix

### The Relationship Between *m* and *τ*

As the model was developed for the phase transition of solids, it has been extensively studied in its thermodynamic context. In the following, we set 1*/τ* = *k*. The rate constant for an isothermal system is given by

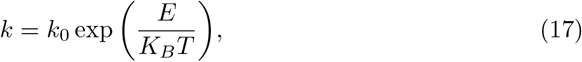

where *K*_*B*_ and *T* denote the Boltzmann constant and the temperature, respectively. *k*_0_ is the pre-exponential factor defined by Arrhenius equation. Lastly *E* represents the overall activation energy, given through

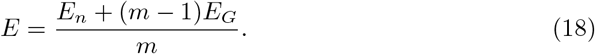

*E*_*n*_ and *E*_*G*_ denote the activation energy for *n* and *G*, respectively. Eq 17 describes a relationship between 1*/τ* = *k* and *m* ([12]). Remarkably, this seems to hold true in the biological context, which provides further support for the validity of the model (S11 Fig). It should be mentioned, though, that missing parameter values for Eq 17 were manually set. A thorough optimisation method was not applied. It hence can only provide some indication.

**S1 Table.** The DC changes for different experimental configurations and data sets.

| Experimental setup | 5 min | 20 min | 60 min | Total |
| --- | --- | --- | --- | --- |
| *TCR* setup: **model** | 0.405 | 0.525 | 0.258 | 0.441 |
| *TCR* setup: **data** | 0.433 | 0.644 | 0.452 | 0.209 |
| *Gene* setup: **model** | 0.226 | 0.396 | 0.216 | 0.241 |
| *Gene* setup: **data** | 0.242 | 0.621 | 0.342 | 0.231 |

**S2 Table.** The number of models per region, before and after applying the requirements for parameter ranges. IGR abbreviates intergenic regions.

| Experimental setup | Region name | #Total | #Filtered |
| --- | --- | --- | --- |
| *Gene* | TS | 4973 | 4356 |
|  | NTS | 4973 | 2294 |
|  | IGR + | 4067 | 1591 |
|  | IGR - | 4067 | 1583 |
| *TCR* | TS start | 1878 | 1865 |
|  | TS centre | 1878 | 1703 |
|  | TS end | 1878 | 1367 |
|  | NTS start | 1878 | 840 |
|  | NTS centre | 1878 | 1080 |
|  | NTS end | 1878 | 1117 |
|  | IGR | 1763 | 650 |

**S3 Table.** Number of non-random relationship between model parameters and sequencing data over *k*. The number of *k*NN models that could find a correlation between model parameters and genomic context. *k* ∈ {5, 10, 20, 50, 100}. We defined a link to be significant if at least three out of five *k* find non-random relationship. - means that data was not used in this setup. NET denotes NET-seq data, ND is nucleosome density, and meres give the relative distance to centromeres or telomeres. Suffices S, C, and E denote start, centre and end of an area. NTCR are non-TCR areas. IGR are intergenic/non-transcribed regions.

|  | *TCR* setup |  |  |  |  |  |  | *Gene* setup |  |  |  |
| --- | --- | --- | --- | --- | --- | --- | --- | --- | --- | --- | --- |
|  | TS S | TS C | TS E | NTS S | NTS C | NTS E | NTCR | TS | NTS | IGR+ | IGR- |
| NET | 5 | 5 | 5 | 0 | 0 | 0 | - | 5 | 3 | - | - |
| Size | 5 | 5 | 5 | 5 | 5 | 5 | - | 5 | 5 | - | - |
| ND | 3 | 1 | 0 | 0 | 0 | 0 | 5 | 5 | 0 | 5 | 5 |
| Abf1 | 0 | 4 | 4 | 0 | 0 | 0 | 5 | 5 | 5 | 0 | 4 |
| H2A.Z | 5 | 5 | 5 | 4 | 4 | 4 | 3 | 5 | 5 | 3 | 0 |
| Meres | 0 | 0 | 0 | 0 | 0 | 0 | 0 | 5 | 0 | 0 | 0 |

**S1 Fig.**
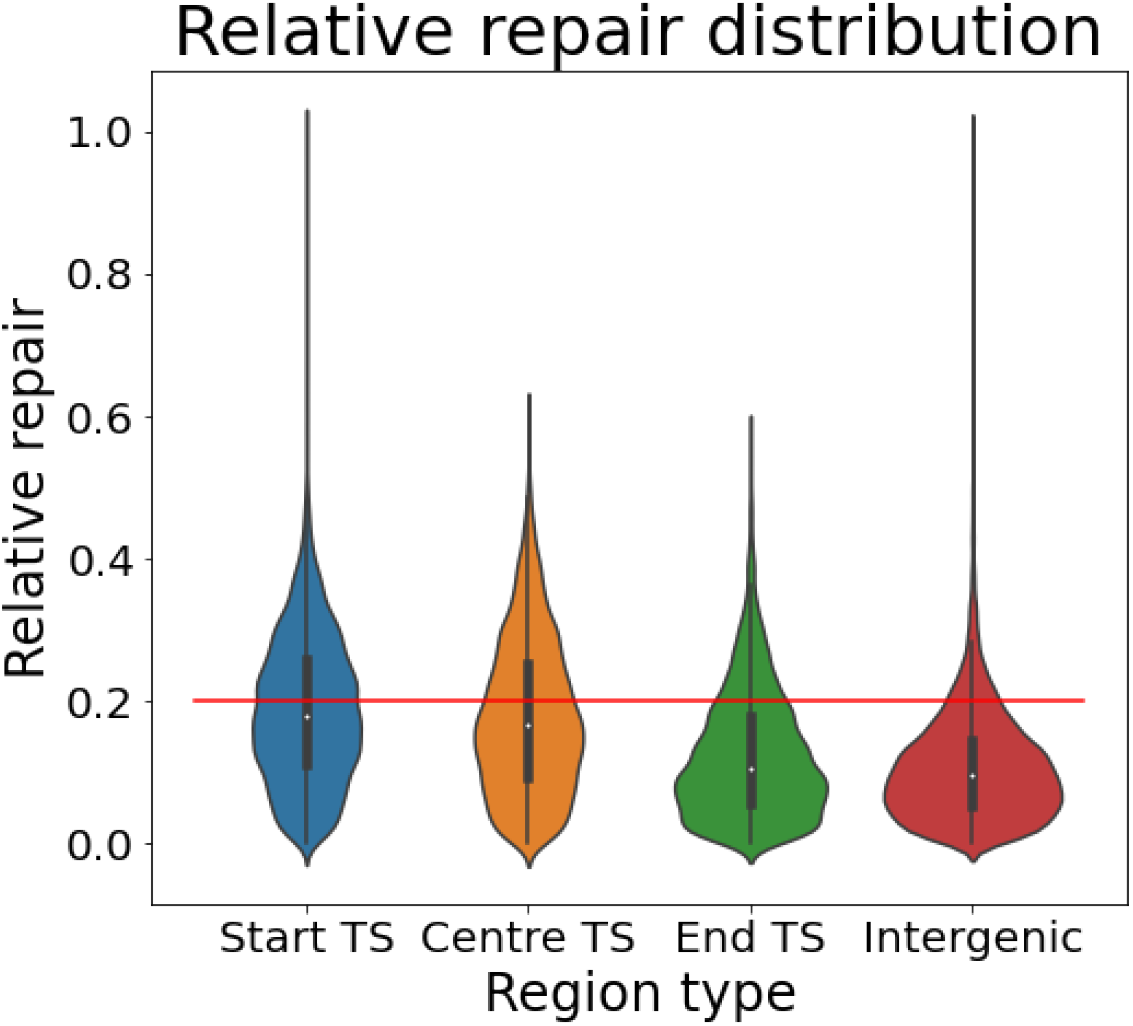
Relative repair distribution over genomic areas. Relative repair in non-transcribed regions is chiefly lower than 20% within the first 20 minutes (88.95%). Genic areas with stronger repair dynamics are thus likely supported by TCR. For all other transcripts, we cannot exclude the possibility that they are exclusively repaired by GGR.

**S2 Fig.**
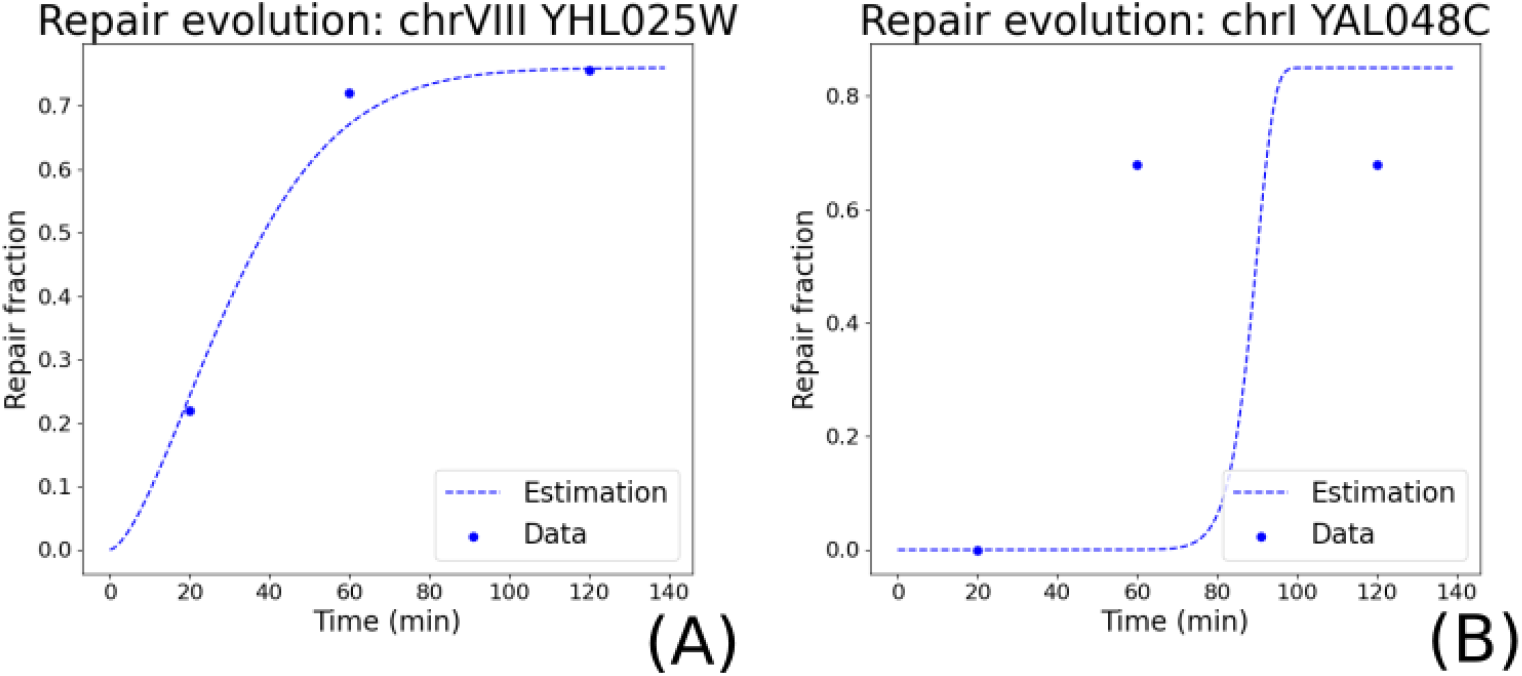
Example for model predictions. (A) The *SNF6* gene can be well approximated. (B) However, *GEM1* exhibits no repair within the first 20 minutes, which results in a switch-like behaviour. Finding an accurate *θ* is difficult.

**S3 Fig.**
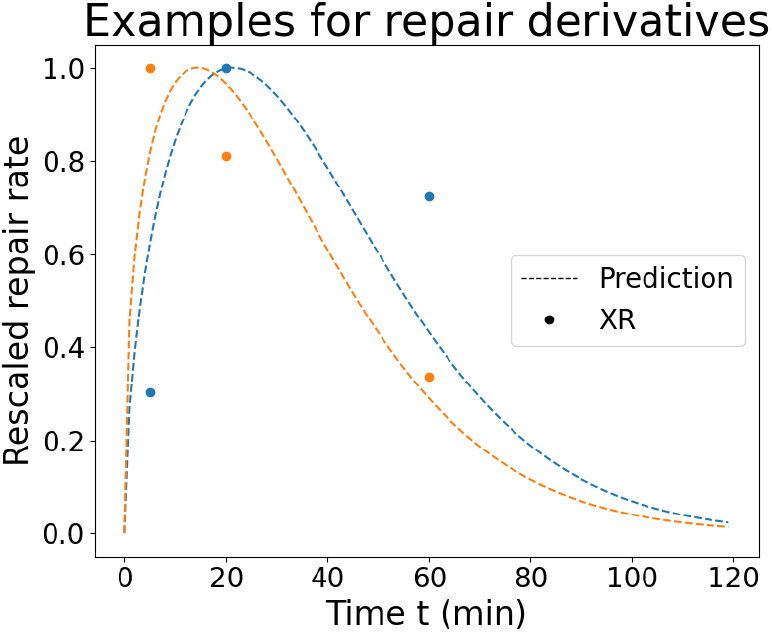
Example for model prediction and XR-seq data over time. When re-scaling XR data and repair rate prediction between 0 and 1, both follow clearly similar trends. The example in blue has its largest XR-seq value at 20 minutes post-radiation, whereas the orange instance shows biggest repair rates after 5 minutes. This is indeed captured by the model.

**S4 Fig.**
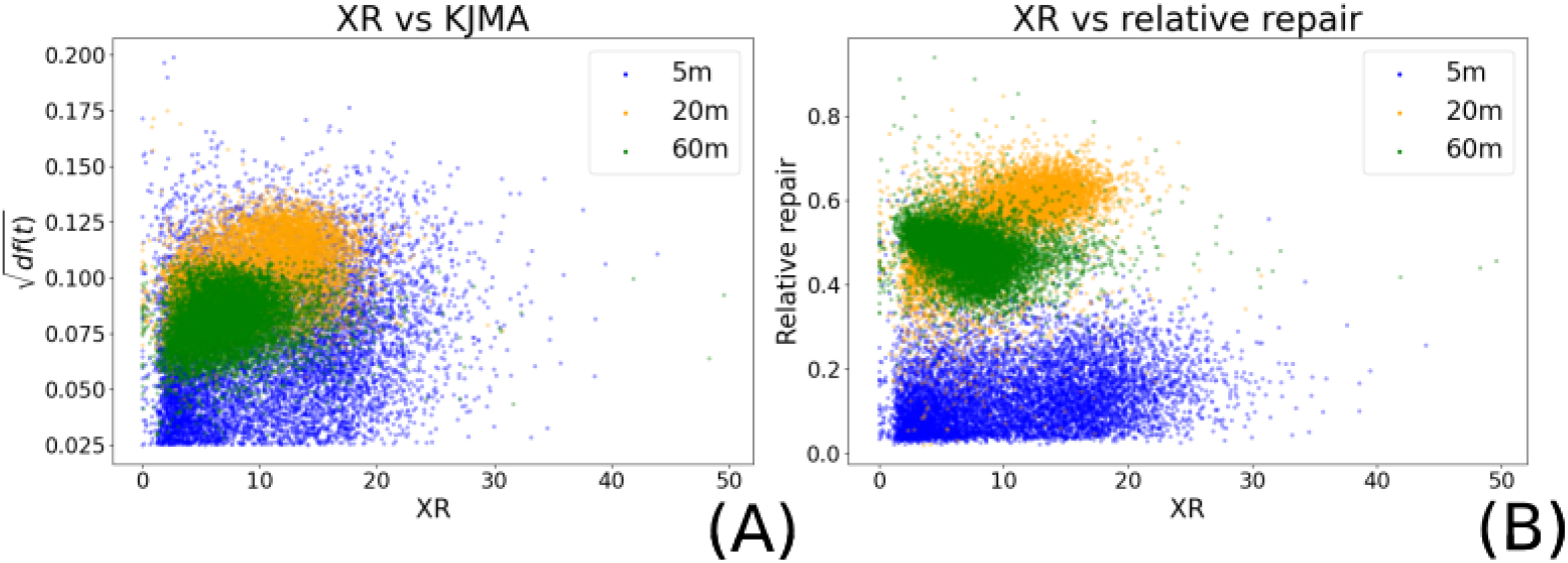
Model predictions with respect to XR-seq data in the *gene* setup. (A) The model predictions in the *gene* configuration are less correlated with the XR-seq data than in the *TCR* setup (DC=0.241). (B) When correlating the *relative repair rate* and the XR-seq data in the *gene* setup, the DC is as low as for the model predictions (DC=0.231). Therefore, we assume that the weak linkage is due to the data segmentation.

**S5 Fig.**
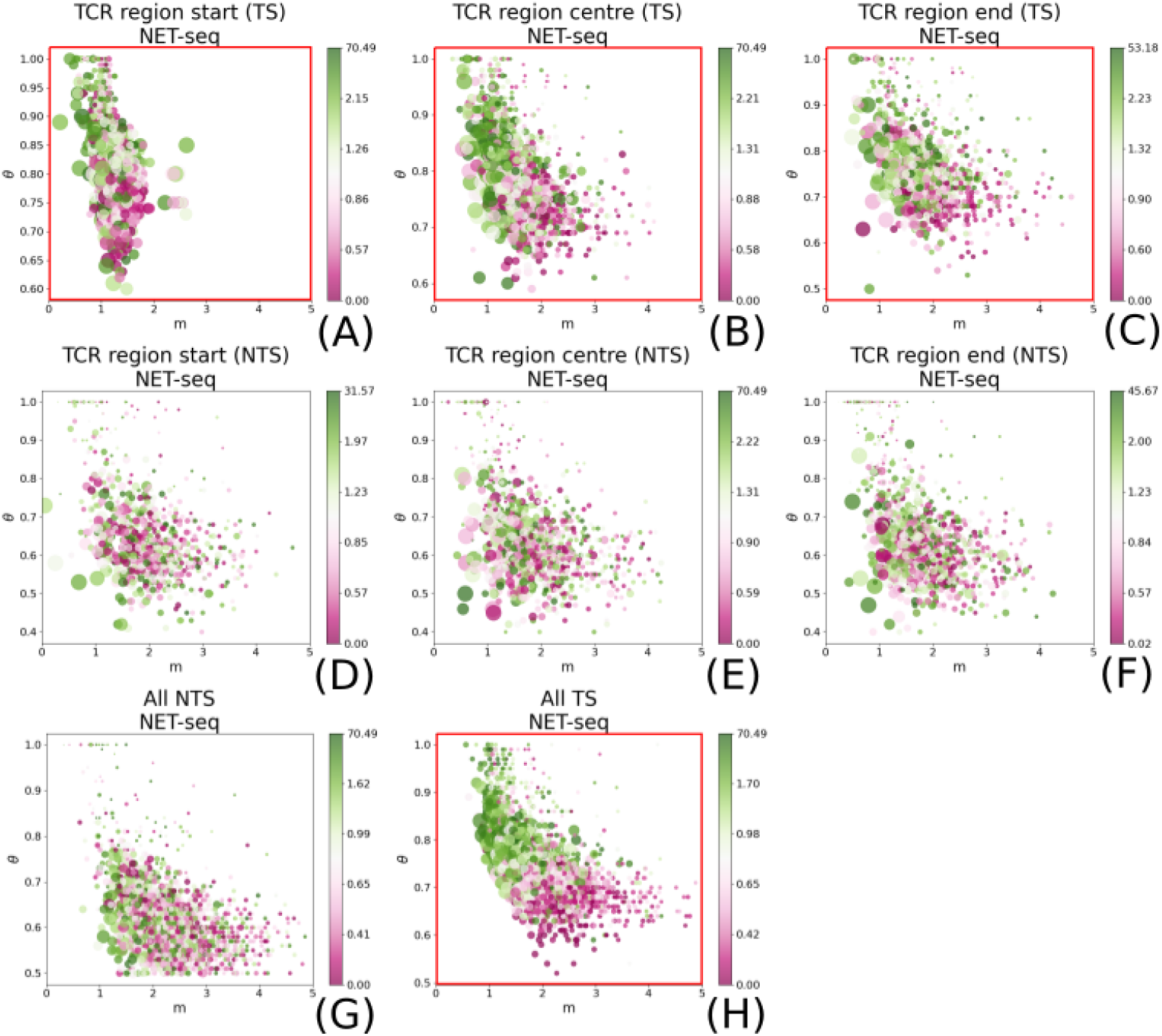
Model parameters with respect to transcription rate. Our results for the transcription rate support the hypothesis that it influences repair on the TS. Red boxes indicate a significant correlation.

**S6 Fig.**
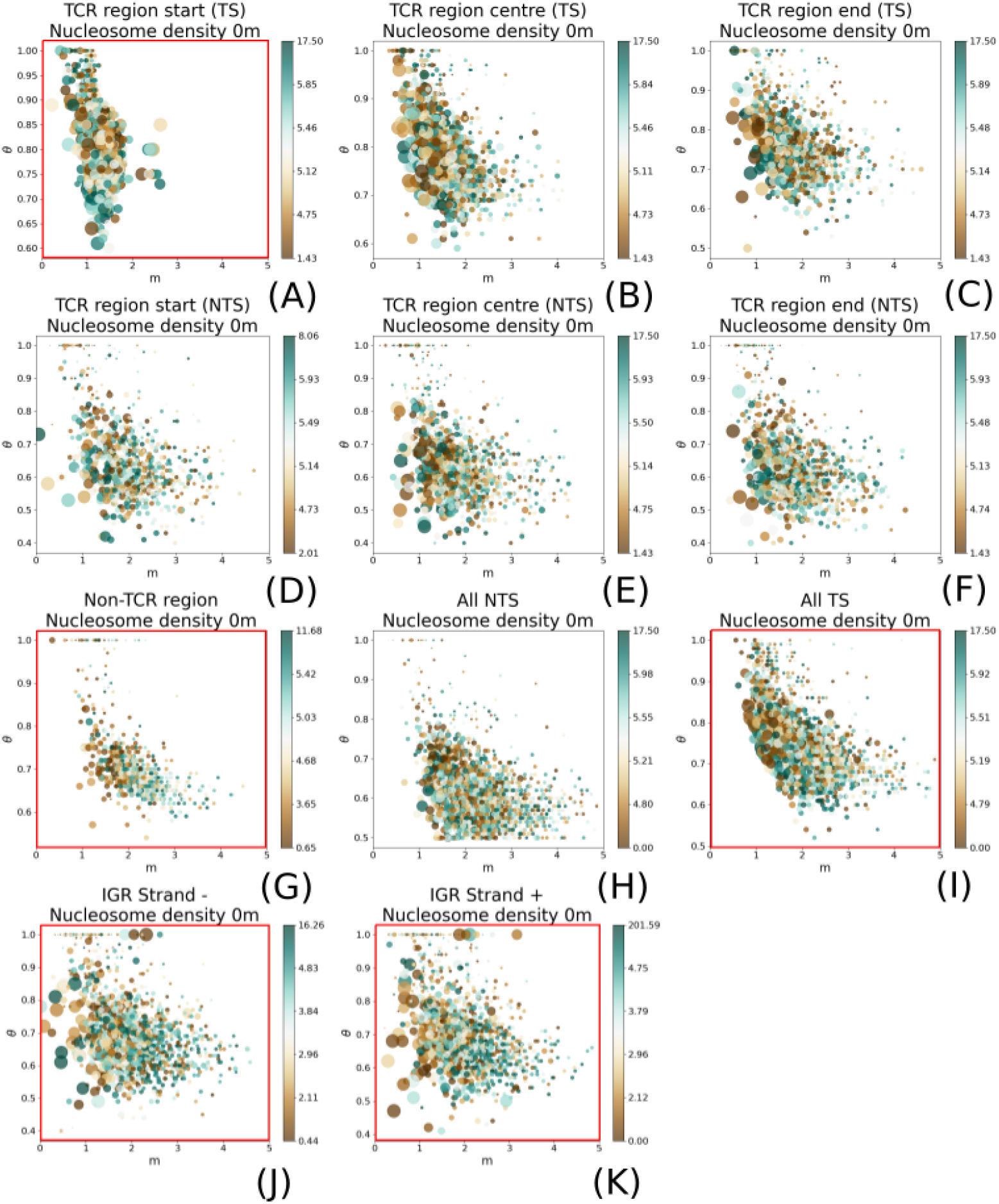
Model parameters with respect to nucleosome density. Nucleosome density is seemingly influencing repair in non-transcribed/non-TCR regions as well as the beginning of the TCR TS and the TS in the *gene* setup. Red boxes indicate a significant correlation.

**S7 Fig.**
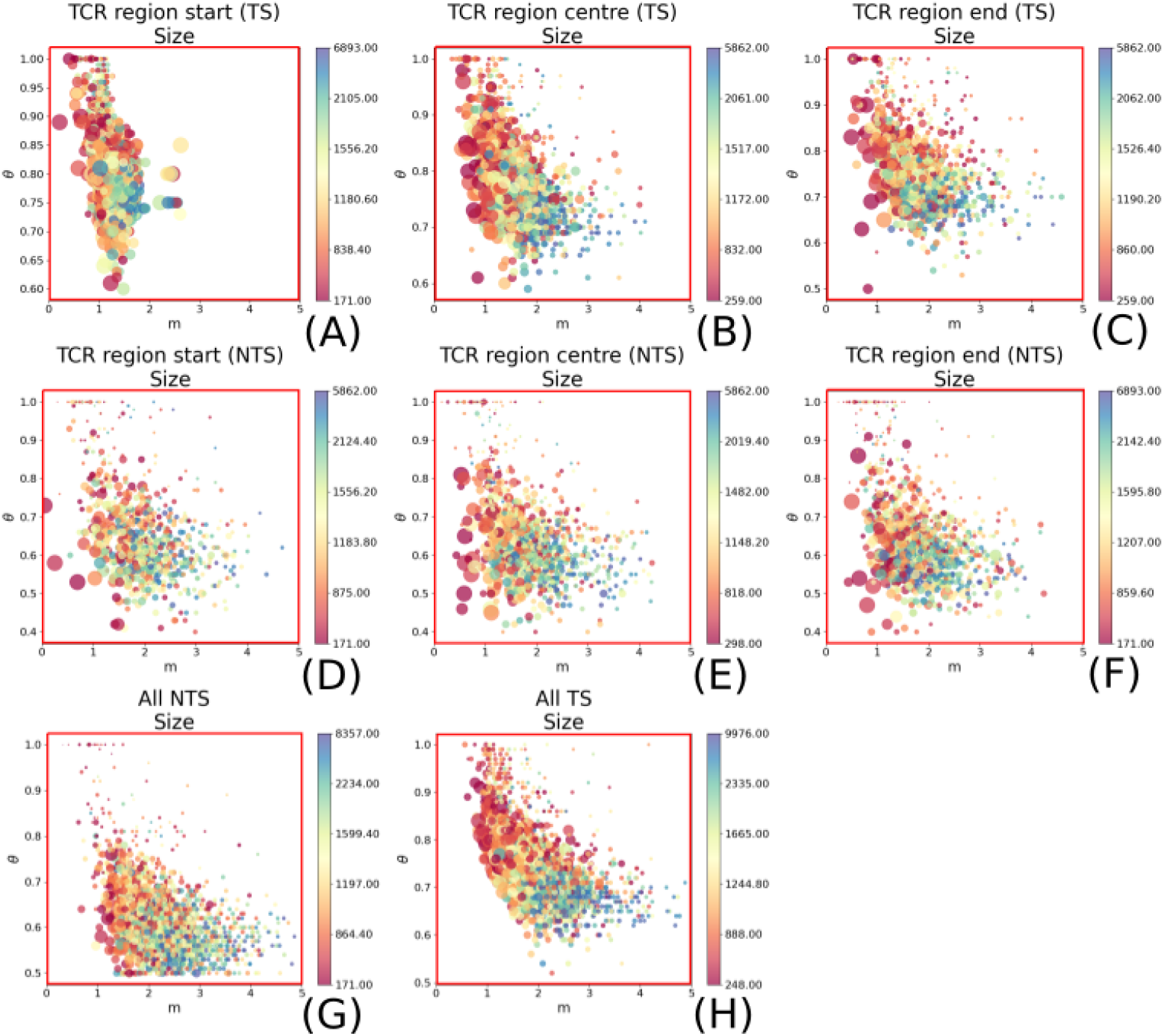
Model parameters with respect to size. The size is clearly influencing repair for both, TS and NTS in the *TCR* and *gene* configuration. Red boxes indicate a significant correlation.

**S8 Fig.**
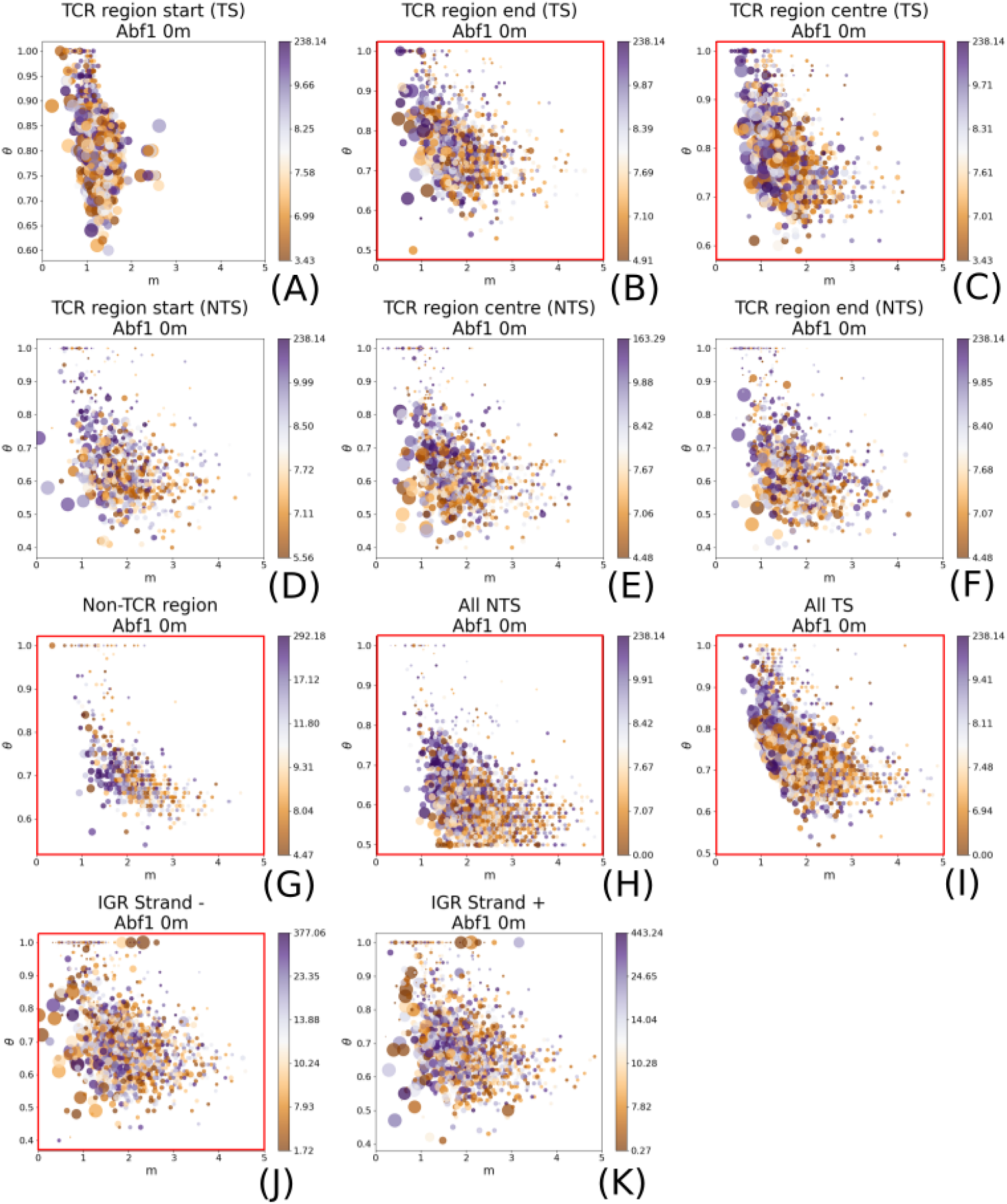
Model parameters with respect to Abf1. Red boxes indicate a significant correlation.

**S9 Fig.**
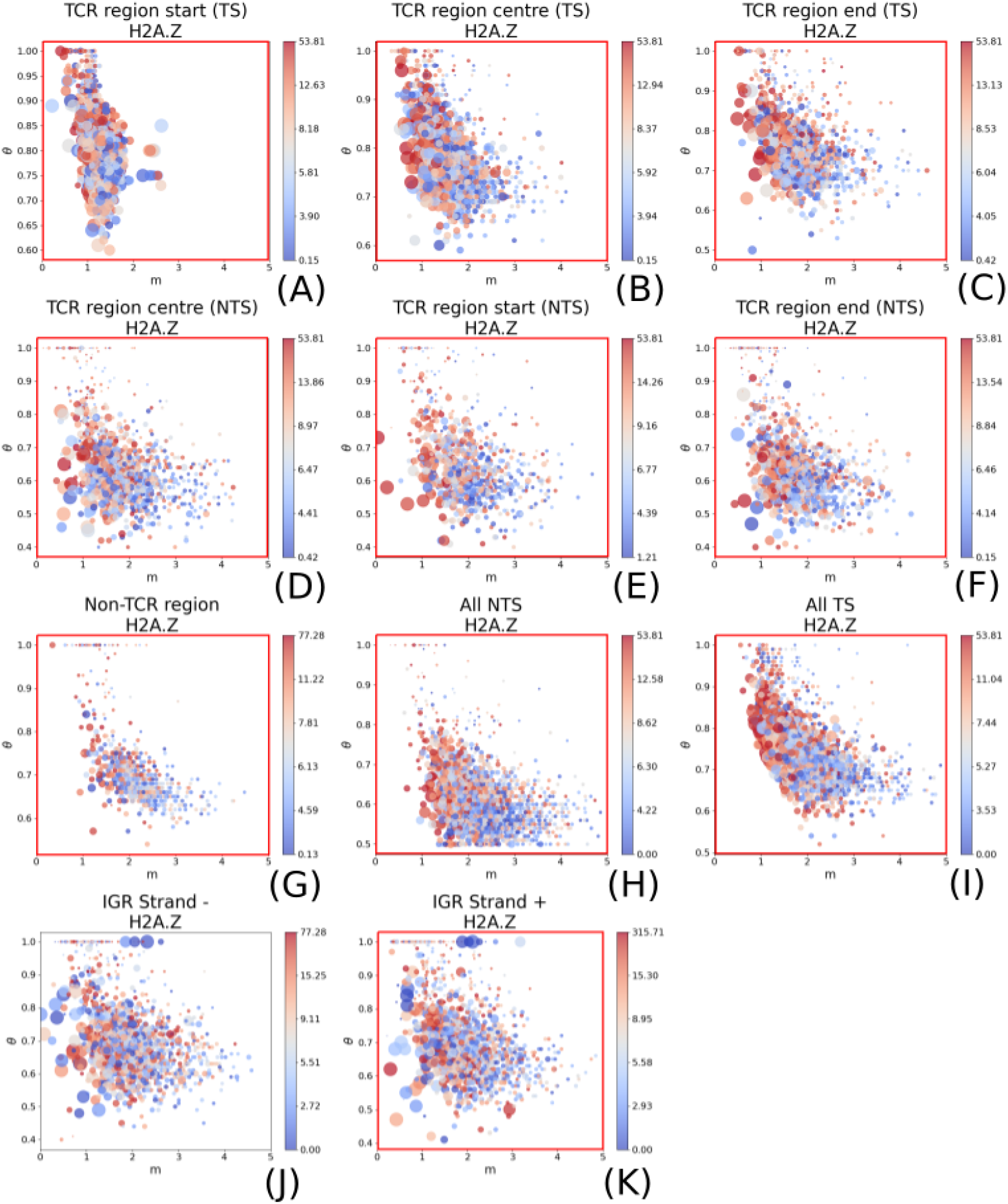
Model parameters with respect to H2A.Z. Red boxes indicate a significant correlation.

**S10 Fig.**
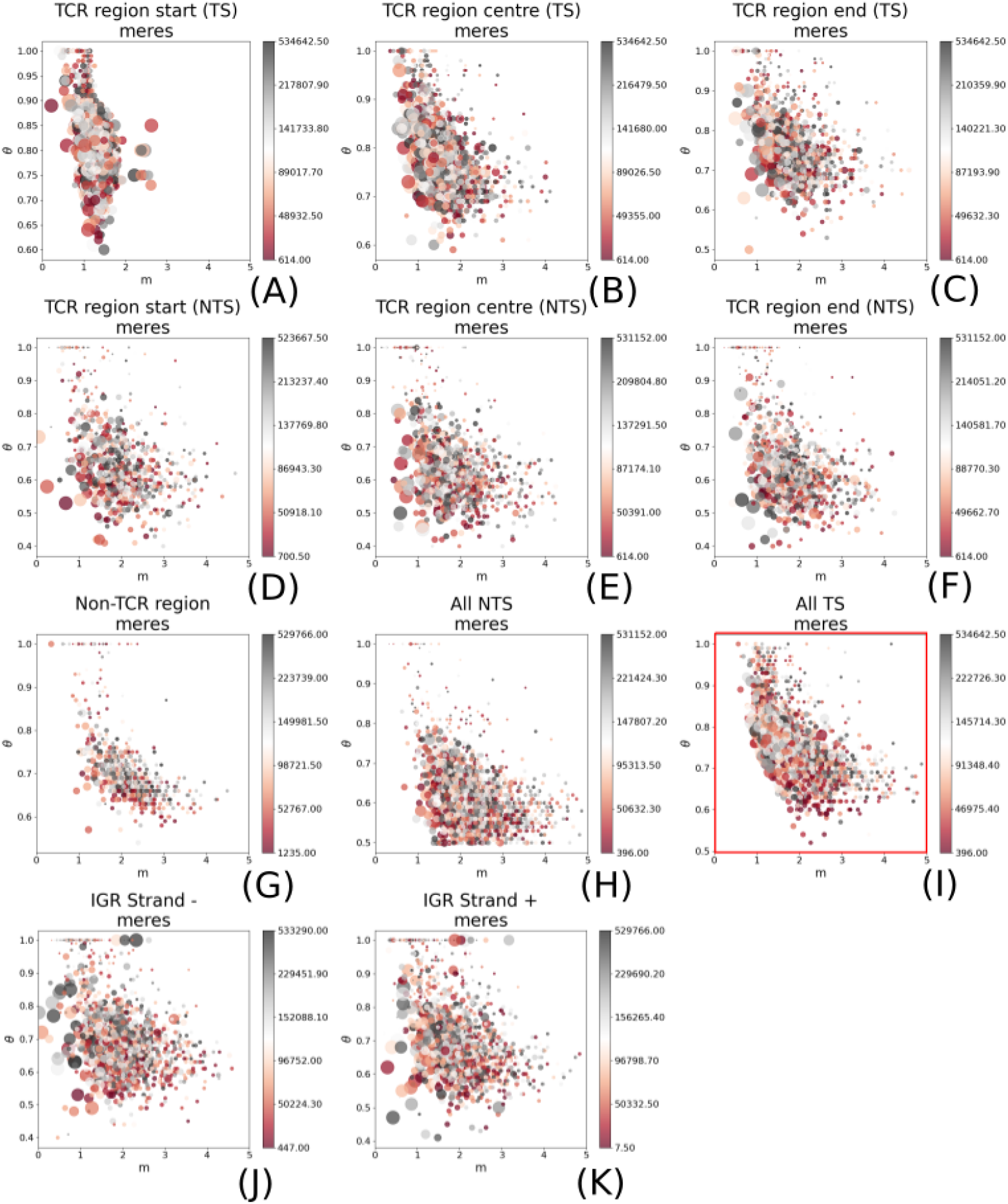
Model parameters with respect to cetromeres and telomeres. With the exception of the TS in the *gene* setup, the distance to telomeres or centromeres (shortened with *meres*) does not affect repair dynamics. Red boxes indicate a significant correlation.

**S11 Fig.**
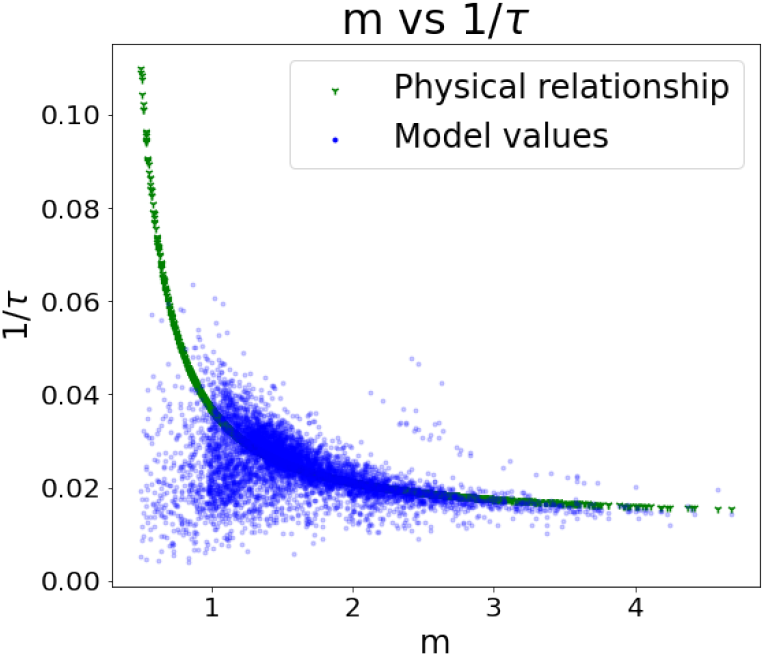
*m* as a function of the turnover rate 1*/τ*. *m* and the turnover rate 1*/τ* follow a trajectory that is predicted by the thermodynamic properties of the model. This provides further support for its applicability (see S6 Appendix).

**S12 Fig.**
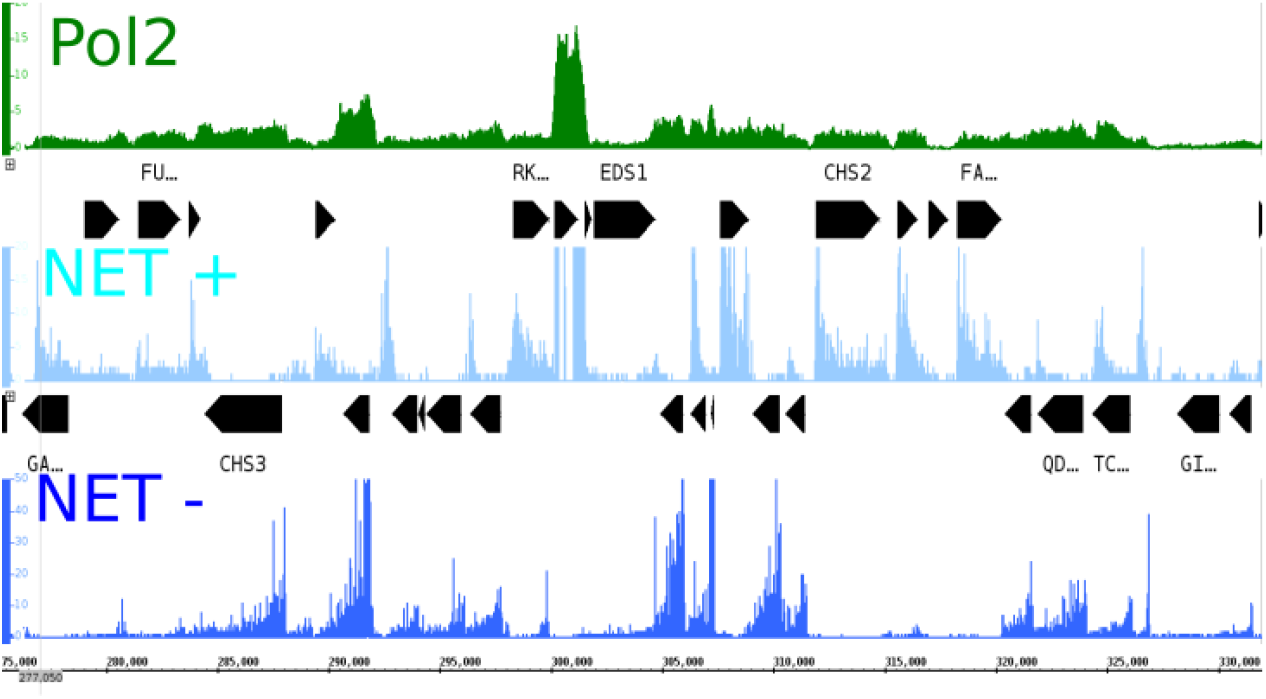
Example of sequencing data representing transcription rate. The example of the NET-seq signal in comparison to the Pol2 ChIP-seq data probed by [13] shows that Pol2 exhibits a constant augmentation of the signal amplitude at transcribed regions, NET-seq data decreases as a function of distance from the TSS. The Pol2 data is coloured in green, whereas NET-seq is given in blue (light blue represents the Watson, and dark blue is the Crick strand). The example is given for chromosome II around *CHS2* and *CHS3*.

**S13 Fig.**
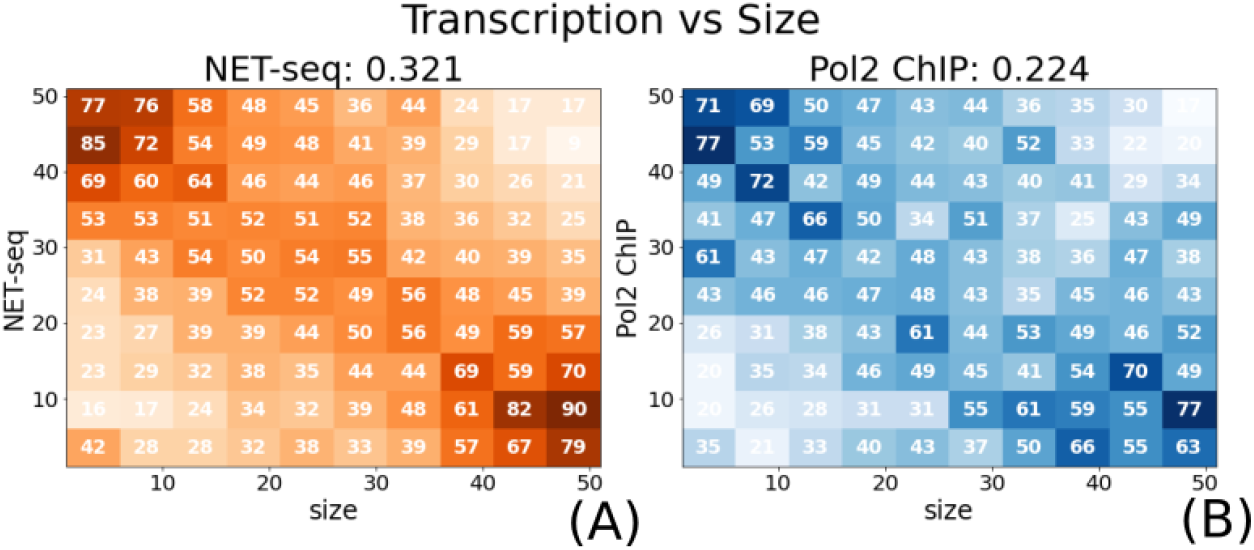
Correlation between size and transcription rate. (A) The histogram distribution of NET-seq transcription with respect to size reveals that smaller genes tend to have higher transcription rates than larger genes. (B) This link is weakened when considering Pol2 ChIP-seq data. Data was discretised into 50 bins such that each contains the same number of transcripts. We use the 95th percentile to remove strong outliers.

**S14 Fig.**
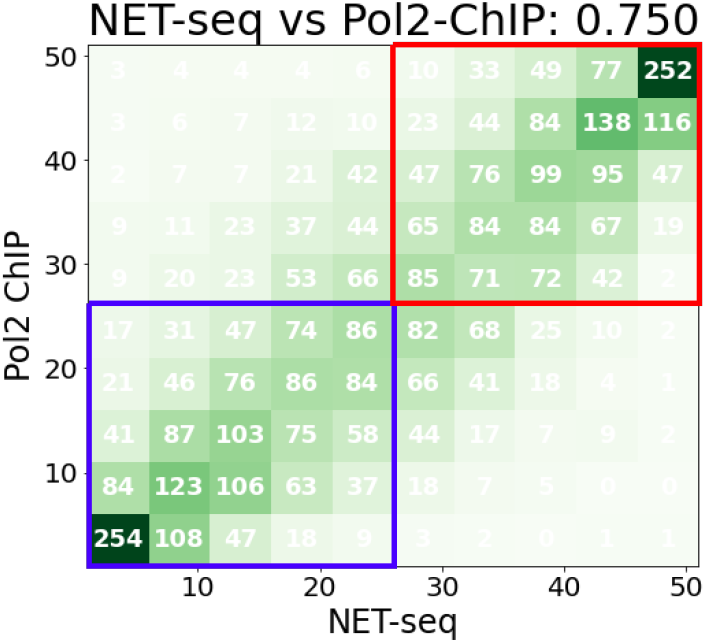
Correlation between NET-seq and Pol2 ChIP-seq data. NET-seq data and Pol2 ChIP-seq signal are strongly related (DC=0.75). As we consider only two groups of genes with respect to transcription, i.e. genes with a low (blue) or high transcription rate (red), we can confirm that the majority of regions fall into the same category.

